# An inconvenient association between granzyme A and Nicotinamide Nucleotide Transhydrogenase

**DOI:** 10.1101/2021.03.16.435730

**Authors:** Daniel J. Rawle, Thuy T. Le, Troy Dumenil, Cameron Bishop, Kexin Yan, Eri Nakayama, Phillip I. Bird, Andreas Suhrbier

## Abstract

Granzyme A (GzmA) is a serine protease secreted by cytotoxic lymphocytes, with GzmA^-/-^ mouse studies informing our understanding of GzmA’s physiological function. We show herein that GzmA^-/-^ mice have a mixed C57BL/6J and C57BL/6N background and retain the full length Nicotinamide Nucleotide Transhydrogenase (*Nnt*) gene, whereas *Nnt* is truncated in C57BL/6J mice. Chikungunya viral arthritis was substantially ameliorated in GzmA^-/-^ mice; however, the presence of *Nnt*, rather than loss of GzmA, was responsible for this phenotype by constraining lymphocyte infiltration. A new CRISPR active site mutant C57BL/6J GzmA^S211A^ mouse provided the first insights into GzmA’s bioactivity free of background issues, with circulating proteolytically active GzmA promoting immune-stimulating and pro-inflammatory signatures. Remarkably, k-mer mining of the Sequence Read Archive illustrated that ≈27% of Run Accessions and ≈38% of Bioprojects listing C57BL/6J as the mouse strain, had *Nnt* sequencing reads inconsistent with a C57BL/6J background. The *Nnt* issue has clearly complicated our understanding of GzmA and may similarly have influenced studies across a broad range of fields.

## INTRODUCTION

Granzyme A (GzmA) is a granule trypsin-like serine protease (trypase) secreted by cytotoxic lymphocytes such as NK cells (Fehniger et al., 2007; Wu et al., 2019), NKT cells (Gordy et al., 2011), and CD8+ cytotoxic T lymphocytes (Suhrbier et al., 1991). The traditional view has been that GzmA is a cytotoxic mediator that is secreted into the immunological synapse, entering the target cell via perforin pores, whereupon certain cytoplasmic proteins are cleaved resulting in the initiation of cell death pathway(s) (Liesche et al., 2018; Martinvalet et al., 2008; Wu et al., 2019; Zhou et al., 2020). A key tool in the quest to understand the physiological role of GzmA has been the use of GzmA^-/-^ mice (Ebnet et al., 1995). For instance, control of viral infections can be compromised in GzmA^-/-^ mice (Loh et al., 2004; Mullbacher et al., 1996; Pereira et al., 2000; Riera et al., 2000), with cytotoxic lymphocytes from these mice reported to be less able to kill target cells (Pardo et al., 2002; Pardo et al., 2004; Shresta et al., 1999; Susanto et al., 2013). Like granzyme B (GzmB), GzmA has thus been classified as a cytotoxic granzyme (Golstein and Griffiths, 2018; Mpande et al., 2018; Muraro et al., 2017; Zhou et al., 2020), although in several studies a role for GzmA in mediating cellular cytotoxicity was not observed (Ebnet et al., 1995; Joeckel and Bird, 2014; Regner et al., 2009; Regner et al., 2011; Smyth et al., 2003). In a range of settings GzmA has also been associated with the promotion of inflammation, providing an additional or alternative view of its physiological role, although consensus on mechanisms has remained elusive (Metkar et al., 2008; Park et al., 2020; Santiago et al., 2020; Santiago et al., 2017; Schanoski et al., 2019; Shimizu et al., 2019; van Daalen et al., 2020; Wensink et al., 2015; Wilson et al., 2017), with a number of potential intracellular and extracellular targets for GzmA reported. These include pro-IL-1β (Hildebrand et al., 2014), SET complex proteins (Mandrup-Poulsen, 2017; Mollah et al., 2017), gasdermin B (Zhou et al., 2020), mitochondrial complex I protein NDUFS3 (Martinvalet et al., 2008), protease activated receptors (Hansen et al., 2005; Sower et al., 1996; Suidan et al., 1994; Suidan et al., 1996), TLR2/4 (van Eck et al., 2017) and TLR9 (Shimizu et al., 2019). GzmA^-/-^ mice have also been used to show a role for GzmA in *inter alia* diabetes (Mollah et al., 2017), cancer (Santiago et al., 2020), bacterial infections (Garcia-Laorden et al., 2016; Garcia-Laorden et al., 2017; van den Boogaard et al., 2016) and arthritis (Santiago et al., 2017). GzmA’s bioactivity has generally (Plasman et al., 2014; Schanoski et al., 2019; Zhou et al., 2020), but not always (Shimizu et al., 2019; van Eck et al., 2017), been associated with GzmA’s protease activity, with circulating GzmA in humans shown to be proteolytically active (Spaeny-Dekking et al., 1998).

After infection with the arthritogenic alphavirus, chikungunya virus (CHIKV) (Suhrbier, 2019), GzmA^-/-^ mice showed a reduced mononuclear cellular infiltrate and substantially reduced overt arthritic foot swelling, when compared to C57BL/6J (6J) mice (Wilson et al., 2017). However, we show herein that active site mutant GzmA^S211A^ mice generated by CRISPR on a 6J background showed no significant differences in CHIKV arthritic foot swelling when compared with 6J mice. The apparent contradiction was resolved when it emerged that GzmA^-/-^ mice had a mixed 6J and C57BL/6N (6N) background. Critically, GzmA^-/-^ mice retained expression of the full length Nicotinamide Nucleotide Transhydrogenase (*Nnt*) gene, whereas 6J mice have a truncated *Nnt* with a 5 exon deletion (Δ*Nnt*8-12) (Fontaine and Davis, 2016). By constraining lymphocyte migration, the presence of full length *Nnt* (rather than absence of GzmA) emerged to be responsible for amelioration of CHIKV arthritic foot swelling in GzmA^-/-^ mice. As much of our understanding of the physiological role of GzmA comes from studies in GzmA^-/-^ mice, we used the GzmA^S211A^ mice to gain new insights into GzmA function that were not compromised by genetic background.

NNT is located in the inner mitochondrial membrane and catalyzes the conversion of NADH plus NADP^+^ to NAD^+^ plus NADPH, while H^+^ is pumped from the inter-membrane space into the mitochondrial matrix (Rydstrom, 2006). NNT thereby sustains mitochondrial anti-oxidant capacity through generation of NADPH (Ward et al., 2020), with the loss of active NNT in 6J mice associated with increased levels of reactive oxygen species (ROS) (McCambridge et al., 2019; Meimaridou et al., 2018; Ronchi et al., 2013; Rydstrom, 2006). As ROS are involved in many cellular processes (Gambhir et al., 2019; Sun et al., 2020), we sought to determine how many other studies might have been affected by the *Nnt* gene differences. K-mer mining of RNA-Seq data sets deposited in the NCBI Sequence Read Archive (SRA) revealed that ≈27% of Run Accessions and ≈38% of Bioprojects listing the mouse strain as “C57BL/6J” had *Nnt* reads inconsistent with a 6J background. Although reported as under-appreciated in the metabolism literature (Fontaine and Davis, 2016), the *Nnt* (and perhaps other background) problems clearly extends well beyond this field and is not restricted to GzmA^-/-^ mice.

## RESULTS

### CHIKV inflammatory arthritis in GzmA^S211A^ mice

We reported previously (Wilson et al., 2017) that the inflammatory arthritis induced by CHIKV infection (manifesting as overt foot swelling) was significantly lower in GzmA^-/-^ mice than in C57BL/6J (6J) mice; an observation we confirm herein (Fig. 1a). Injection of proteolytically active, but not proteolytically inactive recombinant mouse GzmA, induced inflammatory foot swelling, arguing GzmA proinflammatory activity requires it protease activity (Schanoski et al., 2019). To confirm and extend these findings, a new homozygous GzmA active site mutant mouse was generated using CRISPR technology in 6J mice, with the reactive site serine changed to alanine (GzmA^S211A^) (Susanto et al., 2013) (Supplementary Fig. 1a-c). Loss of enzyme activity was confirmed by BLT assays (Suhrbier et al., 1991) (Supplementary Fig. 1d). Intra-cellular staining (Schanoski et al., 2019) showed that expression of the GzmA proteins in resting splenic NK cells (Fehniger et al., 2007) was comparable for GzmA^S211A^ and 6J mice (Supplementary Fig. 1e).

**Fig. 1.**
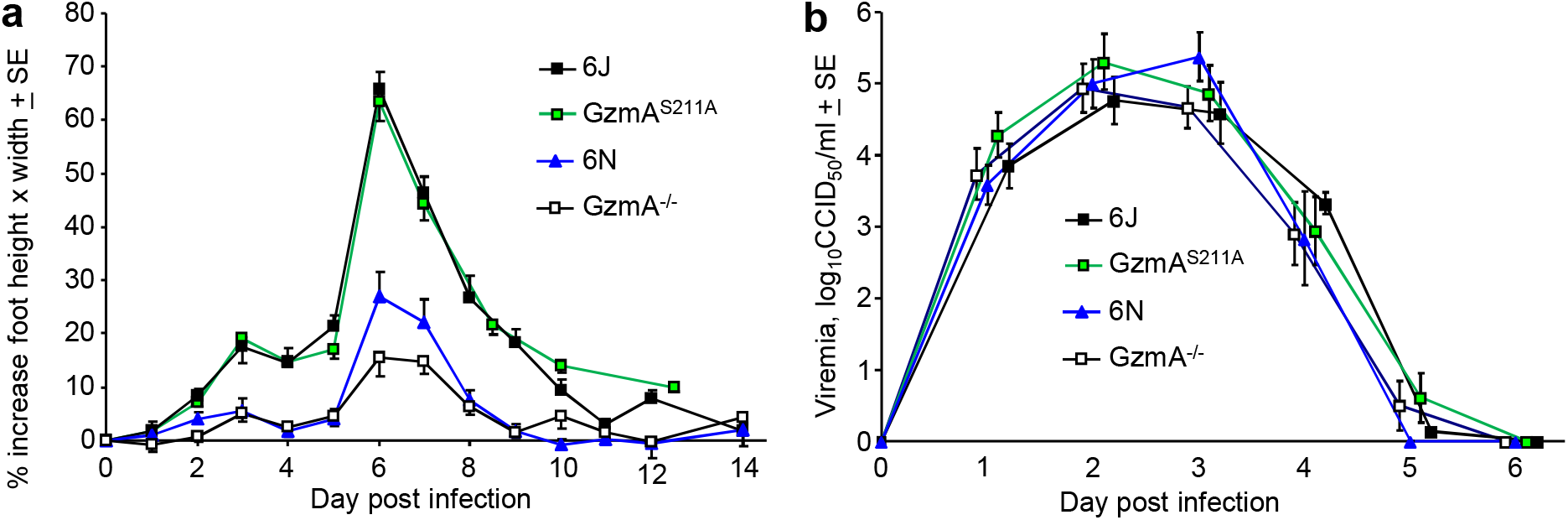
CHIKV infection in GzmA^-/-^, GzmA^S211A^, 6N and 6J mice. **a.** Percent increase in foot swelling for the indicated mouse strains. Data is from 2-4 independent experiments with 5-6 mice (10-12 feet) per group per experiment. From day 3 to day 10, feet from GzmA^S211A^ and 6J mice were significantly more swollen than feet from GzmA^-/-^ and 6N mice (Kolmogorov-Smirnov tests, p<0.002). **b** Viremia for the mice in a (6N n=12, GzmA^S211A^ n=18, GzmA^-/-^ n=15, 6J n=17).

When infected with CHIKV, viral titers in feet were not significantly different for GzmA^S211A^ and 6J mice (Supplementary Fig. 1f), arguing that enzymically active GzmA has no significant antiviral activity against CHIKV. This is consistent with our previous study using GzmA^-/-^ mice that concluded that GzmA has no important antiviral function (Wilson et al., 2017). Surprisingly, however, foot swelling following CHIKV infection was <u>not</u> significantly different between the GzmA^S211A^ and 6J mice (Fig. 1a). If GzmA’s bioactivity (Wilson et al., 2017) depends on its protease activity (Schanoski et al., 2019) the results from GzmA^-/-^ and GzmA^S211A^ mice would appear to provide contradictory results.

The effective amelioration of CHIKV arthritic foot swelling in 6J mice following treatment with Serpinb6b (an inhibitor of GzmA) also supported the view that GzmA promotes inflammation in this setting (Wilson et al., 2017). However, Serpinb6b also inhibited CHIKV foot swelling in GzmA^S211A^ mice (Supplementary Fig. 1g), arguing that Serpinb6b can inhibit other unknown proinflammatory proteases. The contention is supported by the broad inhibitory activity of the human orthologue, SerpinB6 (Strik et al., 2004).

Taken together these new data argued that the role of GzmA in promoting CHIKV arthritis and the results obtained from GzmA^-/-^ mice (Wilson et al., 2017) required some re-evaluation.

### GzmA^-/-^ mice have a mixed C57BL/6N - C57BL/6J background

To reconcile the apparent contradictory data from GzmA^S211A^ and GzmA^-/-^ mice, and cognisant of previously described issues with knock-out mice (Teoh et al., 2014), whole genome sequencing (WGS) of GzmA^-/-^ mice was undertaken (NCBI SRA; PRJNA664888). This analysis unequivocally demonstrated that GzmA^-/-^ mice have a mixed genetic background, with ≈60% of the genome showing single nucleotide polymorphisms (SNPs) and indels present in the C57BL/6N (6N) genome (Mekada et al., 2015; Simon et al., 2013), with the rest reflecting a 6J background (Supplementary Fig. 2). The strain origin of the BL/G-III ES cells used to generate GzmA^-/-^ mice was only reported as C57BL/6 (Ebnet et al., 1995). As ES cells from 6J mice have a low rate of germline transmission, ES from 6N mice cells were frequently used to generate knockout mice (Fontaine and Davis, 2016), with our studies clearly showing that BL/G-III ES cells also have a 6N background. The reported backcrossing of GzmA^-/-^ mice onto C57BL/6 mice (Mullbacher et al., 1999) also clearly did not involve extensive backcrossing onto 6J mice.

Importantly, a large body of literature has resulted from the use of inbred GzmA^-/-^ mice (and GzmA^-/-^/GzmB^-/-^ double knock-out mice derived from them) and used 6J mice as controls, without being aware of the presence and potential confounding influence of the 6N background (Supplementary Table 1).

### GzmA^-/-^ mice and 6N (GzmA^+/+^) mice both have reduced foot swelling after CHIKV infection

The simplest explanation for the different CHIKV foot swelling phenotypes in the two GzmA mutant mouse strains is that a gene from the 6N background is confounding the overt foot swelling phenotype in the GzmA^-/-^ mice. To test this contention, GzmA^-/-^ mice and 6N mice (which are GzmA^+/+^) were infected with CHIKV. 6N mice showed a significant reduction in foot swelling when compared with C57BL/6J mice (Fig. 1a), arguing that the 6N background, rather than deletion of GzmA, was primarily responsible for amelioration of foot swelling in GzmA^-/-^ mice.

There were no significant differences in viremia for 6N, 6J, GzmA^-/-^ or GzmA^S211A^ mice (Fig. 1b), arguing that the reduced foot swelling in 6N and GzmA^-/-^mice was not due to reduced viral loads.

### RNA-Seq of CHIKV foot swelling for GzmA^-/-^ vs 6J mice

To gain insights into the reduced foot swelling in GzmA^-/-^ mice (Fig. 1a), RNA-Seq was undertaken on day 6 (peak arthritis) to compare gene expression in feet from CHIKV-infected GzmA^-/-^ mice versus 6J mice (NCBI Bioproject PRJNA664644; full gene list in Supplementary Table 2a). Differentially expressed genes (DEGs) (n=1073) were generated after application of a q<0.01 filter (Supplementary Table 2b). We have previously undertaken RNA-Seq of CHIKV arthritis in 6J mice (Wilson et al., 2017) (NCBI Bioproject PRJNA431476) and have re-analyzed the fastq files using STAR aligner and the more recent mouse genome build (GRCm38 Gencode vM23); all genes are shown in Supplementary Table 2c and a DEG list (with filters q<0.01, fold change >2) is shown in Supplementary Table 2d. In summary, two gene sets and DEG lists were generated; (i) CHIKV arthritis in GzmA^-/-^ vs CHIKV arthritis in 6J mice, and (ii) CHIKV arthritis in 6J vs naïve 6J mice.

Using Gene Set Enrichment Analysis (GSEA), it emerged that DEGs down-regulated during CHIKV arthritis in GzmA^-/-^ mice (relative to CHIKV arthritis in 6J mice) were not significantly enriched in the up-regulated genes for CHIKV arthritis in 6J mice (relative to mock infection) (Fig. 2a). (Up-regulated DEGs in GzmA^-/-^ mice only reflected reduced epidermal damage, Supplementary Fig. 3). Consistent with Fig. 2a, of the 1557 up-regulated DEGs associated with CHIKV arthritis in 6J mice (Supplementary Table 2d), only a few were down-regulated DEGs for CHIKV arthritis in GzmA^-/-^ mice (Supplementary Table 2b) (Fig. 2b). The reduced foot swelling in GzmA^-/-^ mice (Fig. 1a) was thus <u>not</u> primarily associated with the loss of proinflammatory genes (previous shown to be induced by CHIKV in 6J mice), instead it was associated with the downregulation of a largely distinct set of genes (Fig. 2a, b). In summary, the purported GzmA-dependent component of the CHIKV arthritic response (Wilson et al., 2017) was not lost in GzmA^-/-^ mice, instead arthritis ameliorating in GzmA^-/-^ mice was due to the down-regulation of a distinct and unrelated set of genes.

**Fig. 2.**
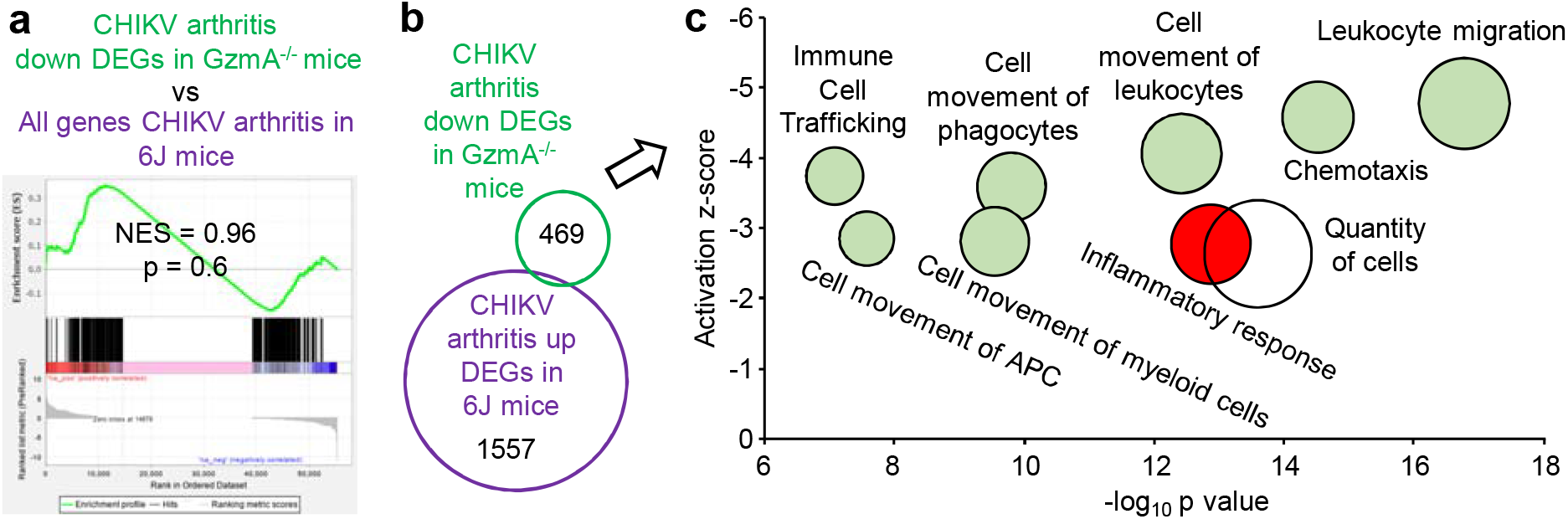
RNA-Seq GzmA^-/-^ vs 6J. **a** GSEA of down-regulated DEGs in feet of CHIKV infected GzmA^-/-^ mice vs feet of CHIKV infected 6J mice (Supplementary Table 2b) versus a full list of genes (pre-ranked by fold change) from feet of CHIKV infected 6J mice vs uninfected 6J mice (Supplementary Table 2c). **b** RNA-Seq identified 1557 DEGs up-regulated in feet of CHIKV infected 6J mice vs uninfected 6J mice (Supplementary Table 2d). RNA-Seq of feet of CHIKV infected 6J mice versus CHIKV infected GzmA^-/-^ mice identified 469 down-regulated DEGs in GzmA^-/-^ mice (Supplementary Table 2b). **c** Selected IPA *Diseases and Function* annotations of the 469 down-regulated DEGs (full set of annotations shown in Supplementary Table 2f).

### Reduced leukocyte migration signatures in GzmA^-/-^ mice

When the 1073 DEGs (CHIKV arthritis in GzmA^-/-^ versus CHIKV arthritis in 6J mice; Supplementary Table 2b) were analyzed for *Diseases and Functions* using Ingenuity Pathway Analysis (IPA), the dominant annotations were associated with decreased cell movement (often leukocyte migration), with a surprising paucity of annotations associated with inflammation (Supplementary Table 2e). When the 469 down-regulated DEGs were used in this analysis, the dominance of decreased cell movement annotations became more apparent, with >40% of these down-regulated DEGs associated with cell movement annotations (Fig. 2c; Supplementary Table 2f). These findings are consistent with immunohistochemistry data showing significantly reduced T cell and NK cells in the arthritic infiltrates in feet after CHIKV infection of GzmA^-/-^ mice (Wilson et al., 2017), with NK cells (Teo et al., 2015), and in particular CD4 T cells, key drivers of alphaviral arthropathy (Nakaya et al., 2012; Suhrbier, 2019; Teo et al., 2013). IPA Up-Stream Regulator (USR) analysis of the 1073 DEGs further illustrated the paucity of inflammation signatures, with the previously identified dominant proinflammatory cytokine signatures associated with CHIKV arthritis (e.g. TNF, IFNγ, IL-1β (Wilson et al., 2017)) not present in the top 100 of the ≈500 down-regulated USRs in GzmA^-/-^ mice (Supplementary Table 2g).

Taken together these results argue that the reduced inflammatory infiltrate (Wilson et al., 2017) and the reduced foot swelling (Fig. 1a) during CHIKV arthritis in GzmA^-/-^ mice had less to do with reduced inflammatory mediator activity, but instead appeared to be associated with a distinct and intrinsic constraint of lymphocyte migration in GzmA^-/-^ mice.

### GzmA^-/-^ mice have an intact Nicotinamide Nucleotide Transhydrogenase gene

Although there are multiple genes associated with inflammation and/or arthritis that differ between GzmA^-/-^ and 6J mice (Supplementary Table 3), *Nnt* was well placed as a candidate for the cell migration phenotype (Supplementary Fig. 2c). The function of NNT is primarily to reduce levels of mitochondrial ROS (McCambridge et al., 2019; Meimaridou et al., 2018; Ronchi et al., 2013; Rydstrom, 2006; Ward et al., 2020) and ROS promotes cell migration in a range of settings (Franchina et al., 2018; Mousslim et al., 2017; Sakai et al., 2018; Tamborindeguy et al., 2018). 6N mice have a full length *Nnt* gene with 21 protein-coding exons, whereas 6J mice have an in-frame 5-exon deletion removing exons 7-11 (Freeman et al., 2006). Confusingly, the MM10 mouse build numbers the *Nnt* exons differently and includes the non-coding exon 1 that is located before the ATG start site. According to this numbering (which is used herein), 6J mice have lost exons 8-12 of 22 exons. The *Nnt* gene is located only ≈6.2 megabases from the GzmA gene on mouse chromosome 13, so >30 backcrosses would be required to segregate these two loci (Silver, 2008) (Supplementary Fig. 4).

Alignment of the WGS of GzmA^-/-^ mice (PRJNA664888) to the standard 6J MM10 mouse genome build allowed identification of the neomycin cassette insertion site into the GzmA gene that was used to generate the GzmA^-/-^ mice (Ebnet et al., 1995) (Fig. 3a). Curiously, this alignment shows a 12 nucleotide insertion in the 6J genome at the *Nnt* exon 8-12 deletion junction (Fig. 3b). The 12 nucleotides are also absent in other 6N WGS data (Supplementary Fig. 5a), indicating this is not a unique feature of GzmA^-/-^ mice. This insertion in 6J may have accompanied deletion of *Nnt* exons 8-12 during the generation of 6J mice (Fontaine and Davis, 2016) (Supplementary Fig. 5b).

**Fig. 3.**
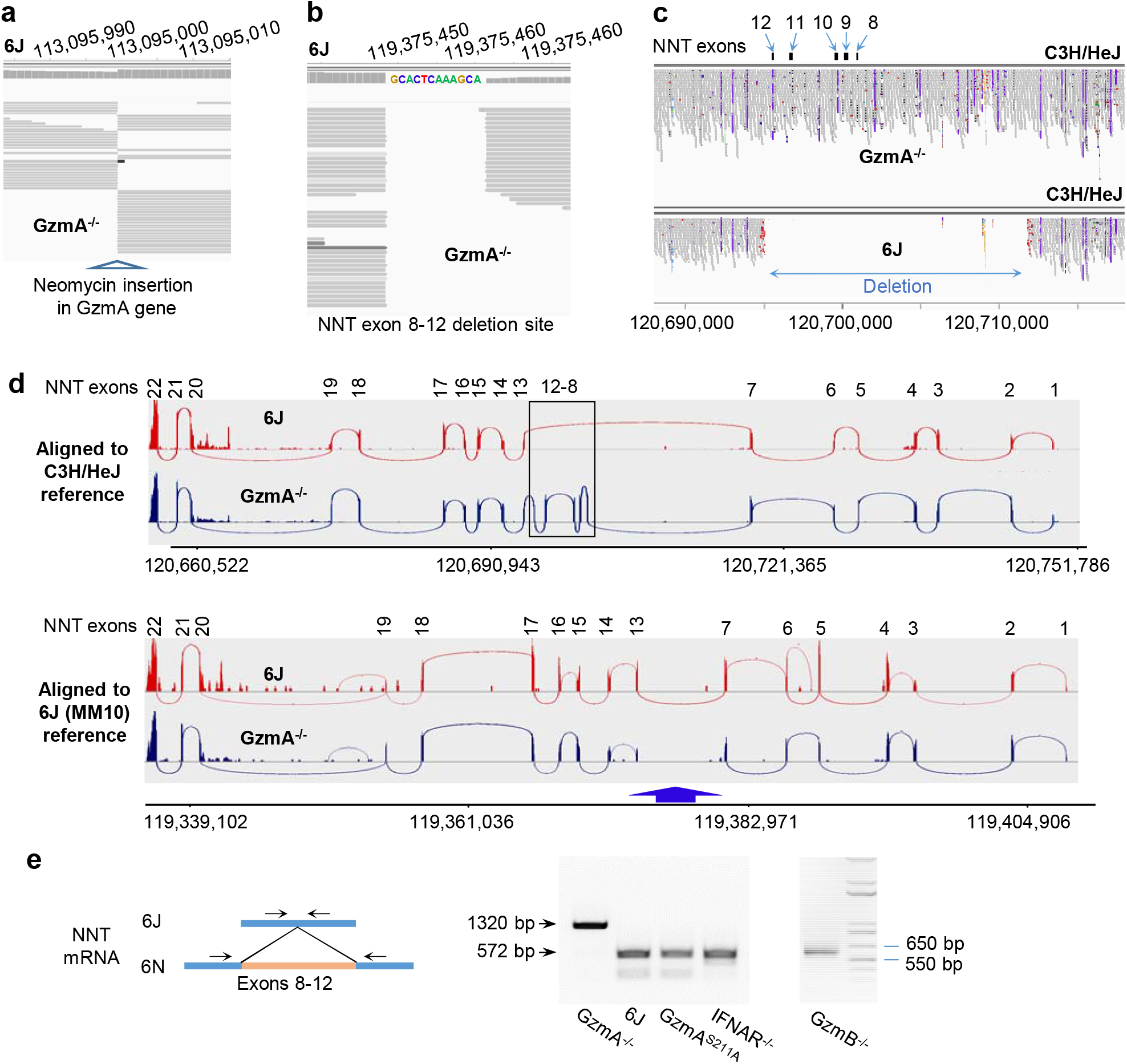
*Nnt* deletion in 6J but not GzmA^-/-^ mice. **a** WGS of GzmA^-/-^ mice aligned to the 6J (MM10) reference genome build, illustrating the insertion site of the neomycin cassette into the GzmA gene to create the knock-out. **b** As for a but showing the site of the *Nnt* deletion, with the additional 12 nucleotides present in the 6J genome. **c** WGS of GzmA^-/-^ and 6J mice aligned to the C3H/HeJ reference genome, illustrating that the *Nnt* deletion present in 6J mice is absent in GzmA^-/-^ mice. The deletion is in chromosome 13; position 120,695,141 to 120,711,874 (C3H/HeJ numbering). **d** Reads from RNA-Seq of CHIKV-infected 6J and GzmA^-/-^ mice aligned to the C3H/HeJ and 6J (MM10) reference genomes showing the Sashimi plot (Integrative Genomics Viewer) for the *Nnt* gene. **e** RT PCR of testes using primers either side of exons 8-12 in the *Nnt* mRNA.

Although a 6N genome sequence is available, it is poorly annotated, hence the C3H/HeJ genome build was used for alignments, as it also has a full length *Nnt* gene (Supplementary Fig. 5c). Alignment of WGS reads from GzmA^-/-^ mice to the C3H/HeJ genome clearly showed that GzmA^-/-^ mice had a full length *Nnt* gene, whereas 6J mice had the expected ≈16 kb deletion (Fig. 3c). The approach (Fig. 3c) was further validated using other 6J and non-6J WGS submissions (Supplementary Fig. 5d).

Sashimi plots of RNA-Seq reads aligned to the C3H/HeJ build clearly illustrated that *Nnt* mRNA from 6J mice was missing exons 8-12, whereas in GzmA^-/-^ mice the full length *Nnt* mRNA was expressed (Fig 3d, top). Alignment to the 6J genome and viewed by Sashimi plot showed that exons 7 and 13 are linked in the *Nnt* mRNA from 6J mice, consistent with expression of a truncated GzmA mRNA species. In contrast, exons 7 and 13 are <u>not</u> linked in the *Nnt* mRNA from GzmA^-/-^ mice (Fig 3d, blue arrow) as the mRNA, but not the MM10 genome build, contains exons 8-12.

These result were confirmed by RT PCR using primers located either side of the exon 8-12 deletion (Huang et al., 2006) (Fig. 3e). GzmA^-/-^ mice have the longer 6N *Nnt* PCR product as this *Nnt* mRNA includes exons 8-12, whereas 6J mice, GzmA^S211A^ (generated by CRISPR on a 6J background), and type I IFN receptor knock-out (IFNAR^-/-^) mice (also on a 6J background (Swann et al., 2007)), all showed a shorter PCR product, consistent with the deletion of exons 8-12 in the *Nnt* mRNA (Fig. 3e). In a separate RT PCR run, GzmB^-/-^ mice (Wilson et al., 2017) were shown to be missing *Nnt* exons 8-12, consistent with a 6J background (Fig. 3e).

### Ameliorated CHIKV arthritic foot swelling in 6N mice is reversed in 6N^Δ*Nnt*8-12^ mice

In order to clearly associate the reduced CHIKV foot swelling in GzmA^-/-^ mice with an intact *Nnt* gene, we generated 6N^Δ*Nnt*8-12^ mice wherein exons 8-12 of *Nnt* were deleted from 6N mice using CRISPR (Supplementary Fig. 6). 6N^Δ*Nnt*8-12^ mice thus have the same *Nnt* exon deletions as 6J mice. As before, CHIKV-induced foot swelling was significantly higher in 6J mice when compared with 6N mice (Fig. 4a). Importantly, foot swelling in 6N^Δ*Nnt*8-12^ mice was significantly higher than in 6N mice, with the *Nnt* exon 8-12 deletion essentially restoring foot swelling levels to those seen in 6J mice (Fig. 4a). There were no significant differences in viremia between the mouse strains (Fig. 4b). This data clearly illustrated that absence of a functional *Nnt* gene plays a major role in the overt foot swelling seen after CHIKV infection of 6J mice. The result also strongly argue that the reduced cell migration (Fig. 2c) and reduced lymphocyte infiltrate (Wilson et al., 2017), and the subsequent reduction in foot swelling (Fig. 1a) seen in GzmA^-/-^ mice (when compared with 6J mice), is primarily due to an intact *Nnt* gene, rather than being due to the loss of GzmA.

**Fig. 4.**
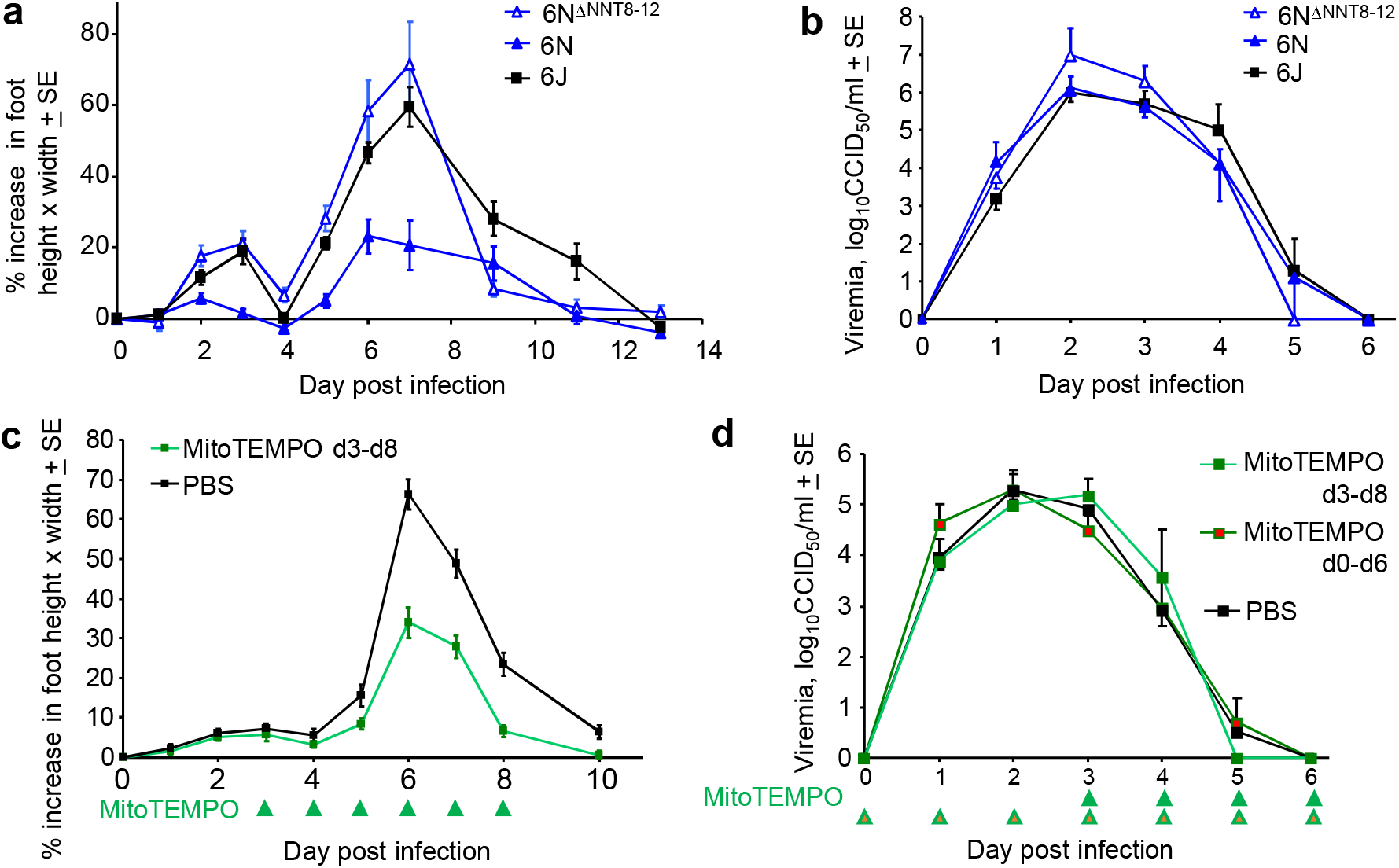
CRIPR 6N^Δ*Nnt*8-12^ mice and MitoTEMPO treatment. **a** 6N^Δ*Nnt*8-12^ mice have the same *Nnt* exon deletions that are present in 6J mice. Age matched female 6N^Δ*Nnt*8-12^, 6N and 6J female mice were infected with CHIKV and foot swelling measured over time (n=5 mice and 10 feet per group). Foot swelling was significantly higher in 6N^Δ*Nnt*8-12^ mice when compared with 6N mice on days 2 through 7 (day 2 p=0.0026, day 7 p=0.0027, t tests, parametric data distributions; days 3-6 p=0.003, Kolmogorov-Smirnov tests, non-parametric data distributions). Foot swelling was significantly lower in 6N mice when compared with 6J mice (day 2 p=0.042, day 6 p=0.001, day 7 p=0.0005, t tests, parametric data distributions; day 3 and 5, p=0.002, Kolmogorov-Smirnov tests, nonparametric data distributions). **b** Viremia for the same mice as in a. **c** Mice were infected with CHIKV and then treated with MitoTEMPO or PBS i.v. daily on days 3-8 post-infection and foot swelling measured (n=5/6 mice and 10/12 feet per group). Statistics by t tests, day 6, 7 and 8 p<0.001. **d** Viremia for the mice in c, with an additional group of 6 mice treated daily with MitoTEMPO from day 0 till day 6.

### Elevated ROS signatures in GzmA^-/-^ mice

As NNT’s primary function is scavenging of mitochondrially generated ROS (McCambridge et al., 2019; Meimaridou et al., 2018; Ronchi et al., 2013; Rydstrom, 2006; Ward et al., 2020), the data argues that the reduced arthritic foot swelling in GzmA^-/-^ mice is due to lower ROS levels. This contention is supported by IPA analysis of DEGs from RNA-Seq data comparing day 6 feet for CHIKV-infected GzmA^-/-^ versus CHIKV-infected 6J mice, where many of the top USRs with negative z-scores were associated with ROS activity; e.g. TGFβ (Liu and Desai, 2015), VEGF (Reichard and Asosingh, 2019), SOX7 (Klomp et al., 2020) and SP1 (Marinho et al., 2014) (Supplementary Table 2g).

A previous publication reported that NNT is a key regulator of adrenal redox homeostasis and provided RNA-Seq analysis of adrenals from *Nnt^-/-^* versus *Nnt^+/+^* mice (Meimaridou et al., 2018). Using Gene Set Enrichment Analysis (GSEA) we show that the DEGs up-regulated in adrenals in *Nnt^-/-^* versus *Nnt^+/+^* mice (Meimaridou et al., 2018) were significantly enriched in genes down-regulated in GzmA^-/-^ (*Nnt^+/+^*) versus 6J (*Nnt^-/-^*) mice (Supplementary Fig. 7a-c). Furthermore, IPA analysis of the core enriched genes from this GSEA identified some of the same ROS-associated USR annotations (Supplementary Fig. 7d) that were found previously for GzmA^-/-^ mice (Supplementary Table 2g). This analysis supports the view that ROS activity is reduced in GzmA^-/-^ mice, and also points to a fundamental role for *Nnt*, with common ROS-associated pathways found in resting adrenals and CHIKV infected feet of *Nnt^-/-^* mice.

### MitoTEMPO ameliorates CHIKV arthritis in 6J mice

MitoTEMPO has been widely used as an experimental anti-oxidant treatment to scavenge mitochondrial ROS in a variety of disease settings (Aoyama et al., 2012; Li et al., 2018; To et al., 2020; Vincent et al., 2020; Wu et al., 2020). Treatment of CHIKV arthritis in 6J mice with MitoTEMPO from day 3 to 8 post infection significantly ameliorated peak foot swelling (Fig. 4c). Viremia was not significantly affected by MitoTEMPO treatment, even when treatment was initiated on day 0 (Fig. 4d). These data support the view that a major contributor to the CHIKV-induced arthritic infiltrate and foot swelling in 6J mice is the excessive generation of mitochondrial ROS, a feature consistent with loss of NNT activity.

### Re-investigation of the physiological role of GzmA using poly(I:C)

Our current understanding of the physiological function of GzmA comes, to a large extent, from multiple studies comparing GzmA^-/-^ (and GzmA^-/-^/GzmB^-/-^) mice with 6J mice (Supplementary Table 1). Many of the reported phenotypes are likely to have arisen, at least in part, from the 6N background, complicating any conclusions regarding the physiological role of GzmA.

To gain new insights into GzmA’s activity *in vivo*, we sought to find an experimental setting where high levels of GzmA are secreted. We have shown previously that humans, non-human primates and mice have elevated serum GzmA levels after infection with CHIKV (Schanoski et al., 2019; Wilson et al., 2017). Infection of mice with a series of other RNA virus also resulted in high serum GzmA levels early in infection, with NK cells identified as the likely source (Schanoski et al., 2019). Resting NK cells constitutively contain abundant levels of GzmA protein (Fehniger et al., 2007), which is usually stored in granules as a mature protease, with the low pH of the granule preventing (premature) proteolytic activity (Stewart et al., 2012).

Polyinosinic:polycytidylic acid (poly(I:C)) can often mimic aspects of the innate responses to RNA viruses (Prow et al., 2017). We thus injected poly(I:C) i.v. into 6J mice and showed that serum GzmA levels reached mean peak serum levels of ≈20 ng/ml of serum after 2 hours, with levels dropping to baseline after 24 hours (Fig. 5a). Although poly(I:C) treatment is known to activate NK cells (Djeu et al., 1979; Fehniger et al., 2007; Miyake et al., 2009; Ngoi et al., 2008), this rapid and prodigious poly(I:C)-induced release of GzmA into the circulation, to the best of our knowledge, has hitherto not been reported. Type I interferons (IFN) are also rapidly induced by poly(I:C) (Dempoya et al., 2012; Santiago-Raber et al., 2003), and NK cells express the type I IFN receptor and can respond to type I IFNs (Madera et al., 2016; Mizutani et al., 2012). Injection of poly(I:C) i.v. into IFNAR^-/-^ mice resulted in a significantly blunted GzmA response (Fig. 5a). Type I IFNs thus appear to augment this rapid GzmA release; however, the absence of type I IFN signaling does not prevent GzmA secretion, consistent with previous data (Schanoski et al., 2019). Poly(I:C) treatment of GzmA^S211A^ mice resulted in serum GzmA levels not significantly different from those seen in 6J mice (Fig. 5a), illustrating that the active site mutation does not significantly affect production, secretion or stability. Finally, although GzmB and GzmA are often considered to be co-expressed (Supplementary Table 1), in this setting no serum GzmB was detected (Supplementary Fig. 8). GzmB and perforin proteins are not expressed in resting NK cells and appear only after ≈24 hours of appropriate stimulation (Fehniger et al., 2007).

### RNA-Seq after poly(I:C) injection in GzmA^S211A^ versus 6J mice

GzmA^S211A^ and 6J mice are both on a 6J genetic background, are both missing *Nnt* exons 8-12, and both show similar levels of serum GzmA secretion after poly(I:C) treatment (Fig. 5a). Thus the only difference between these strains is that GzmA from GzmA^S211A^ mice is enzymically inactive (Supplementary Fig. 1d). GzmA^S211A^ and 6J mice were injected with poly(I:C) (as in Fig. 5a), with feet and spleens removed 6 hours later and analyzed by RNA-Seq (NCBI Bioproject PRJNA666748). The rationale for this time point was to capture early transcriptional events after the peak of serum GzmA. The sample preparation strategy and RNA-Seq data overview is provided in Supplementary Fig. 9. The full gene list for feet is provided in Supplementary Table 4a, with the 199 DEGs listed in Supplementary Table 4b. For spleen the full gene list is shown in Supplementary Table 4c and the 4 DEGs in Supplementary Table 4d. This represents the first study of GMO mice targeting GzmA that is free from the potentially confounding influence of the mixed 6J/6N background.

### Immune/inflammation signatures stimulated by circulating proteolytically active GzmA

Circulating GzmA would appear to have limited influence in the spleen, as only 4 DEGs were identified in the spleen of poly(I:C) treated GzmA^S211A^ mice (Supplementary Table 4d). Interestingly, the only significantly up-regulated DEG in GzmA^S211A^ spleens was Mid1, a gene involved in controlling granule exocytosis by cytotoxic lymphocytes (Boding et al., 2014; Boding et al., 2015). GSEA analyses also illustrated that neither up nor down-regulated DEGs identified in the feet were enriched in spleen (Supplementary Fig. 10), arguing that in this setting the activity of GzmA in the periphery is not significantly recapitulated in spleen.

Of the 199 DEGs identified in feet of poly(I:C) treated GzmA^S211A^ mice, 100 were down-regulated, with the top annotations associated with negative regulation of protease activity and negative regulation of cell signaling after Cytoscape analysis (Fig. 5b; Table 1; Supplementary Table 4e). This result supports the view that GzmA is proteolytically active *in vivo* (Spaeny-Dekking et al., 2000) and that GzmA’s proteolytic activity mediates cell signaling events under physiological conditions. IPA *Disease and Functions* analysis of the 199 DEGs from feet (GzmA^S211A^ versus 6J; Supplementary Table 4b) identified down-regulation (negative z-scores) of a series of inflammation and leukocyte activation signatures (Fig. 5b, Supplementary Table 4f). IPA USR analysis (core analysis with direct and indirect interactions) indicated down-regulation of a series of cytokine, immune receptor and transcription factor USRs (Fig. 5b, Supplementary Table 4g). An IPA USR analysis using the direct only interaction option, which largely limits the analysis to transcription factors, showed STAT6, STAT3, NFATC2, JUN, NFKB1 and STAT1 to be the dominant down-regulated transcription factor signatures in GzmA^S211A^ mice by z-score and p values (Fig. 5b, Supplementary Fig. 11a). These transcription factors are associated with various innate and adaptive immune responses, with NFATC2 playing a central role in activation of T cells during the development of an immune response.

**Table 1.** Cytoscape analysis of down-regulated DEGs GzmA^S211A^ mice. RNA-Seq of feet taken from GzmA^S211A^ vs 6J six hours after poly(I:C) injection provided 199 DEGs, of which 100 were down-regulated in GzmA^S211A^ mice (Supplementary Table 4d). When analyzed by Cytoscape the top annotations were associated with negative regulation (underlined) of protease activities (red) or negative regulation of protein metabolism (which includes anabolism and catabolism). Also significant were a series of annotations associated with negative regulation of cell signaling (yellow). The complete list is shown in Supplementary Table 4e; top annotations are shown here with duplicates removed.

| Category | Description | FDR value |
| --- | --- | --- |
| GO Process | <u>negative</u> regulation of cellular protein metabolic process | 1.91E-06 |
| UniProt Keywords | <u>Protease inhibitor</u> | 9.20E-06 |
| SMART Domains | <u>SERine Proteinase INhibitors</u> | 2.60E-05 |
| GO Process | <u>negative</u> regulation of catalytic activity | 6.73E-05 |
| GO Process | <u>negative</u> regulation of nitrogen compound metabolic process | 7.86E-05 |
| GO Process | <u>negative</u> regulation of cellular metabolic process | 7.86E-05 |
| InterPro Domains | <u>Serpin superfamily</u> | 1.10E-04 |
| GO Process | <u>negative</u> regulation of molecular function | 1.30E-04 |
| GO Function | <u>enzyme inhibitor activity</u> | 1.50E-04 |
| UniProt Keywords | <u>Serine protease inhibitor</u> | 1.50E-04 |
| GO Process | <u>negative</u> regulation of macromolecule metabolic process | 1.90E-04 |
| GO Process | <u>negative</u> regulation of protein modification process | 2.60E-04 |
| GO Process | <u>negative</u> regulation of hydrolase activity | 2.80E-04 |
| GO Function | <u>serine-type endopeptidase inhibitor activity</u> | 6.60E-04 |
| GO Process | <u>negative</u> regulation of phosphate metabolic process | 7.40E-04 |
| GO Function | <u>endopeptidase inhibitor activity</u> | 8.50E-04 |
| GO Process | <u>negative</u> regulation of endopeptidase activity | 0.001 |
| GO Process | <u>negative</u> regulation of intracellular signal transduction | 0.0014 |
| GO Process | <u>negative</u> regulation of protein phosphorylation | 0.0014 |
| GO Process | regulation of protein metabolic process | 0.0015 |
| GO Process | <u>negative</u> regulation of MAPK cascade | 0.0061 |
| GO Process | <u>negative</u> regulation of cellular process | 0.0063 |
| GO Process | <u>negative</u> regulation of biological process | 0.0071 |

STAT3 and NF-κB have previously been shown to be activated in macrophages by recombinant GzmA *in vitro* (Santiago et al., 2020), with monocytes/macrophages reported as targets for GzmA activity in a variety of settings (Garzon-Tituana et al., 2021; Metkar et al., 2008; Santiago et al., 2017; Spencer et al., 2013; Uranga et al., 2016). Interrogation of the Molecular Signature Database (MSigDB) (Subramanian et al., 2005) also found genesets associated with activated monocytes (GSE19888) significantly enriched in the down-regulated genes for GzmA^S211A^ vs 6J mice by preranked (fold change) GSEAs (Supplementary Fig. 11b). This observation supports the contention that monocytes/macrophages are activated by circulating GzmA (Fig. 5b). IPA analysis of the 150 core enriched genes from these GSEAs, also identified STAT6, STAT3, NFATC2, JUN, NFKB1 and STAT1 as significant USRs, even though only 10 of these 150 genes were significant DEGs for GzmA^S211A^ vs 6J (Supplementary Table 4h). These USR signatures would thus appear to be a consistent feature of the GzmA^S211A^ vs 6J RNA-Seq data.

### GzmA’s bioactivity after CHIKV infection and poly(I:C) treatment

The data so far (summarized in Fig. 6a) argues that the presence of functional NNT, rather than the absence of GzmA, causes the reduction in overt day 6 CHIKV arthritic foot swelling (Fig. 1a), with NNT constraining lymphocyte migration (Wilson et al., 2017) (Fig. 2c) by reducing ROS levels (McCambridge et al., 2019; Meimaridou et al., 2018; Ronchi et al., 2013; Rydstrom, 2006) (Supplementary Table 2g). As GzmA’s proteolytic function would appear to be required for its bioactivity (Schanoski et al., 2019) (Fig. 5b; Table 1), the data from CHIKV-infected GzmA^S211A^ mice (Fig. 1a) argues that GzmA is not the major driver of the CHIKV arthritic foot swelling.

**Fig. 5.**
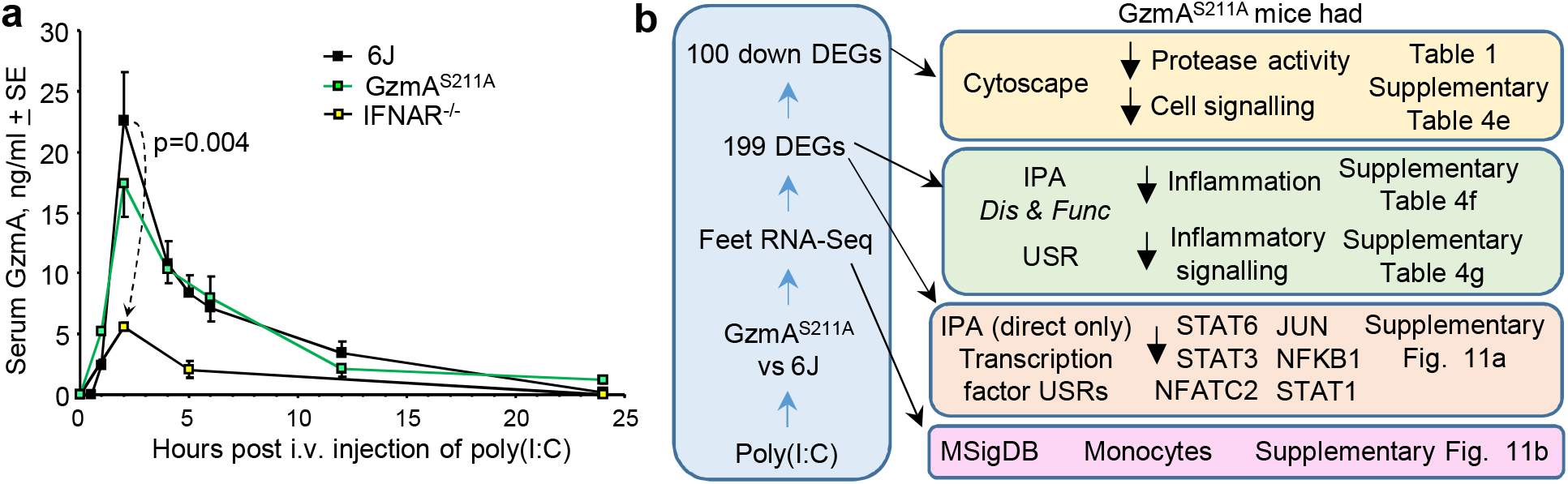
Poly(I:C) injection into GzmA^S211A^ and 6J mice. **a** GzmA^S211A^ and 6J mice were injected i.v. with 250 μg of poly(I:C) in 150 μl of PBS and serum samples were taken at the indicated times assayed for GzmA concentration using a capture ELISA kit (6J n=5-8, IFNAR^-/-^ n=5-6 and GzmA^S211A^ n=3 mice per time point). **b** GzmA^S211A^ and 6J mice were injected i.v. with 250 μg of poly(I:C) and feet removed 6 h later and analyzed by RNA-Seq. The DEGs (Supplementary Table 4b) were analyzed by Cytoscape and IPA. The full gene list (Supplementary Table 4a) was analyzed using the Molecular Signature Database (MSigDB).

We show herein that poly(I:C) induces high levels of circulating GzmA and that type I IFNs potentiate GzmA release (Fig. 6b), with NK cells the likely source of GzmA (Schanoski et al., 2019). RNA-Seq analyses comparing GzmA^S211A^ with 6J mice support the view that circulating proteolytically active GzmA (Table 1) promotes certain immune-stimulating and proinflammatory activities (Schanoski et al., 2019; Shimizu et al., 2019; van Daalen et al., 2020; Wensink et al., 2015), with STAT6, STAT3, NFATC2, JUN, NFKB1 and STAT1 identified as dominant USRs (Fig. 6b. Supplementary Fig. 11a). Consensus regarding the molecular target(s) of extracellular GzmA’s protease actrivity (Fig. 6b, indicated as ??) remains elusive, but may include protease activated receptors (Hansen et al., 2005; Sower et al., 1996; Suidan et al., 1994; Suidan et al., 1996) and/or pro-IL-1 (Hildebrand et al., 2014); the latter can become extracellular when cells lyse (Afonina et al., 2015).

**Fig. 6.**
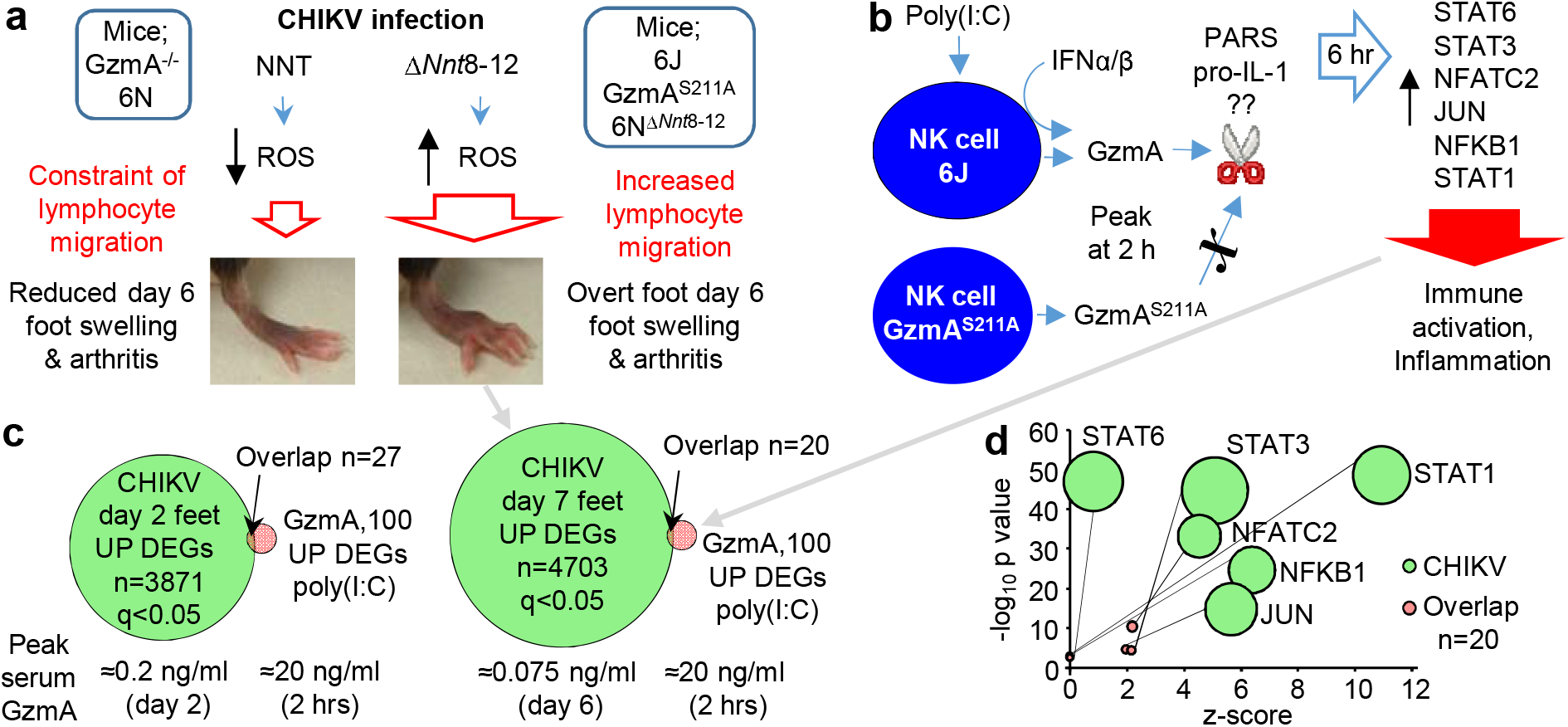
GzmA’s bioactivity after CHIKV infection and poly(I:C) treatment. **a** For CHIKV infection, a primary driver of arthritis and overt foot swelling is loss of functional NNT, with 6J, GzmA^S211A^, and 6N^Δ*Nnt*8-12^ mice all missing exons 8-12 of *Nnt* and showing overt foot swelling. In contrast, mice with an intact *Nnt* gene (GzmA^-/-^ and 6N) show reduced arthritic foot swelling and GzmA^-/-^ mice have been shown to have significantly reduced lymphocyte infiltrates. An important function of NNT is reducing the level of mitochondrially-derived reactive oxygen species (ROS). **b** Using poly(I:C) to induce high levels of GzmA secretion from NK cells, studies in GzmA^S211A^ mice illustrated that proteolytically active circulating GzmA promotes certain immune-stimulating/pro-inflammatory responses. No clear consensus has emerged regarding the molecular target(s) of GzmA (??); two potential extracellular candidates are shown; protease activated receptors and pro-IL-1. **c** DEGs up-regulated in feet by CHIKV infection of 6J mice on day 2 and 7 (applying a filter of q<0.05 to the gene sets in Supplementary Table 2c) were compared with the DEGs up-regulated in feet by GzmA in 6J mice after poly(I:C) treatment (i.e. down-regulated in GzmA^S211A^ mice; Supplementary Table 4b). Overlapping genes (n=27 and 20) are listed in Supplementary Table 4i. The mean peak levels of serum GzmA for each group are shown below the Venn diagrams. **d** The 20 overlapping genes for day 7 were subjected to IPA USR analysis (direct only) and the same USRs shown in b were returned as significant (Supplementary Table 4i). When DEGs up-regulated in feet on day 7 by CHIKV infection (Supplementary Table 2c; applying a q<0.05 filter) were subjected to the same IPA USR analysis, these USRs (amongst many others) were also identified.

GzmA mRNA is significantly elevated in feet during CHIKV infection (Wilson et al., 2017), and serum protein levels peak at ≈0.2 ng/ml on day 2, with a second peak of ≈0.075 ng/ml on day 6 post infection (Schanoski et al., 2019). If GzmA is present and has immune-stimulating and proinflammatory activities, why does it have no significant role in driving the overt CHIKV arthritic foot swelling? Firstly, the peak serum GzmA levels after CHIKV infection @0.2 and ≈0.075 ng/ml) were substantially lower than those seen after poly(I:C) treatment @20 ng/ml) (Fig. 6c), and very much lower than the ≈5 μg of recombinant GzmA injected intraplantar to generate overt foot swelling in the absence of any other stimuli (Schanoski et al., 2019). Secondly, when DEGs up-regulated in feet during CHIKV infection in 6J mice (Supplementary Table 2c, q<0.05), where compared with DEGs up-regulated in 6J mice by GzmA (vs GzmA^S211A^ mice; Supplementary Table 4b), only small overlaps were evident (27 genes for day 2 and 20 genes for day 6) (Fig. 6c). Although the comparisons might be viewed as purely illustrative given the different settings, the small overlap nevertheless argues that the contribution of GzmA to the overall CHIKV arthritic responses is minimal; an observation consistent with GzmA^S211A^ foot swelling data in Fig. 1a. When the 20 overlapping genes for day 6 arthritis (Fig. 6c) were subjected to IPA USR (direct only) analysis, STAT6, STAT3, NFATC2, JUN, NFKB1 and STAT1 annotations remained significant (Supplementary Table 4i). These USRs (amongst many others) are also identified by the same analysis of DEGs up-regulated during CHIKV arthritis (Supplementary Table 2c, q<0.05); however, z-scores were much higher and p values much lower (Fig. 6d). So CHIKV arthritis and GzmA stimulate overlapping immune/inflammation pathways, but GzmA only plays a minor role in stimulating these pathways in CHIKV arthritis.

### 6J SRA accessions with *Nnt* exon reads inconsistent with a 6J background

Given the data presented herein and elsewhere (Bourdi et al., 2011; Fontaine and Davis, 2016; Ripoll et al., 2012; Vozenilek et al., 2018) and given ROS are involved in many cellular processes (Gambhir et al., 2019; Sun et al., 2020), *Nnt* emerges as a legitimate focus of concern (Fontaine and Davis, 2016; Leskov et al., 2017; McCambridge et al., 2019; Mekada et al., 2009; Rydstrom, 2006). We undertook a k-mer mining approach to interrogate the NCBI SRA (Supplementary Fig. 12a), which (at the time of analysis) contained 61,443 RNA-Seq Run Accessions listing “C57BL/6J” in the strain field of the metadata.

For “C57BL/6J” Run Accessions, k-mer mining was used to count the number of RNA-Seq reads with sequence homology to *Nnt* exon 2 or exon 9, with these two exons being of similar length (203 bp for exon 2 and 192 bp for exon 9). RNA-Seq analysis of 6J tissues would ordinarily provide reads for exon 2, whereas the presence of exon 9 reads would be inconsistent with a 6J background. A conservative k-mer mining approach was used; (i) only an exact match for “C57BL/6J” in the strain field was allowed, (ii) Run Accessions with small or large compressed file sizes (<200 Mb and >1500 Mb) were excluded, (iii) nucleotide mismatches for the 31 nucleotide k-mers were disallowed, (iv) where there were technical replicates, only one was mined and (v) a read count of ≥10 per exon was used as a cut-off. This k-mer mining analysis revealed that 1008 Run Accessions had reads aligning to both *Nnt* exons 2 and 9, indicating full length *Nnt* (*Nnt+*), which is not consistent with 6J (Supplementary Table 5a). In contrast, 2469 had exons 2 reads, but no exon 9 reads indicating truncated *Nnt* (*Nnt-*), which is not consistent with 6J (Supplementary Table 5b). Lastly, 267 Run Accessions had equivocal results (Supplementary Table 5c). Therefore ≈27% (1008 of 3744) of Run Accessions listing “C57BL/6J” in the strain field had sequencing reads not consistent with a 6J background. The k-mer mining approach was validated for a selected group of Run Accessions using NCBI BLAST alignments, which illustrated excellent concordance with the k-mer mining read count data (Supplementary Table 5d).

The startlingly high percentage (≈27%) of SRA Accessions listing “C57BL/6J” but having *Nnt* reads inconsistent with 6J, argues that the *Nnt* issue is widely under-appreciated in a broad range of research areas. It should be noted that a large number of Run Accessions (n=206,586) list “C57BL/6” in the strain field and thus do not provide information on the substrain (Mekada et al., 2009) being used.

### Bioprojects comparing mice with truncated *Nnt* to mice with full length *Nnt*

Based on the results of k-mer mining of “C57BL/6J” Run Accessions, Bioprojects (n=373) were grouped into three categories (i) Bioprojects where all the Run Accessions with “C57BL/6J” in the strain field had RNA-Seq reads that were consistent with 6J (*Nnt-*) (62%), (ii) Bioprojects where all the Run Accessions with “C57BL/6J” in the strain field were not consistent with 6J (*Nnt+*) (23%) and (iii) Bioprojects where some Run Accessions with “C57BL/6J” in the strain field were *Nnt-* and others were *Nnt+* (n=57; 15%). Thus 38% (15% plus 23%) of Bioprojects had Run Accessions with “C57BL/6J” strain listings not compatible with a 6J background.

Of the 57 aforementioned Bioprojects, 43 had at least one publication associated with the study. These Bioprojects were then manually interrogated to identify studies where it was clear (from the paper and the metadata) that comparisons had been made between two groups, where all the Run Accessions in one group were *Nnt+*, and all the Run Accessions in the other group were *Nnt-* (Fig. 7; Supplementary Fig. 12b; Supplementary Table 5d and e). Aside from the CHIKV Bioproject described herein, several others emerged (Fig. 7). For example, Mir31^-/-^ mice showed reduced CD8 T cell dysfunction during chronic viral infection when compared to 6J mice (Moffett et al., 2017); however, Mir31^-/-^ mice were *Nnt+* (Fig. 7, Bioproject PRJNA385694; Supplementary Table 5d). Rel^-/-^Nfkb1^-/-^CD4CreRela^fx/fx^ mice were compared with 6J mice to implicate RIPK1 and IKK in thymocyte survival (Webb et al., 2019); however, the control 6J mice were *Nnt+* (Fig. 7, Bioproject PRJEB30085; Supplementary Table 5d). Bruce4 ES cells were reported to be on a 6J background (Ank et al., 2008); however, Bruce4 cells were *Nnt+* (Fig. 7, Accession SRR923434; Supplementary Table 5d). The reported differences between Bruce4 and 6J genomes were thus likely more to do with the background than with genetic instability (Hughes et al., 2007). IL-28RA^-/-^ mice were generated using Bruce4 cells and were compared with 6J mice (Ank et al., 2008); however, RNA-Seq analysis of *IL-28RA^-/-^* mice showed that (like Bruce4 cells) these mice had full length *Nnt* (Supplementary Fig. 13). MyD88^-/-^Trif^-/-^ double knock-out mice were compared with 6J mice during infection with *S. aureus* (Scumpia et al., 2017); however, the 6J (wild-type) mice were *Nnt+* (Fig. 7, Bioproject PRJNA382450; Supplementary Table 5e). Female Cystatin C^-/-^ mice display significantly lower clinical signs of Experimental Autoimmune Encephalomyelitis (EAE) when compared with 6J mice (Hoghooghi et al., 2020); however, Cystatin C^-/-^ mice were all *Nnt+* (Fig. 7, Bioproject PRJNA662247; Supplementary Table 5d). Deletion of *Nr1h3* resulted in reduced chromatin access at a large fraction of Kupffer-cell specific enhancers (Sakai et al., 2019); however, Nr1h3^-/-^, but not the wild-type control, were all *Nnt+* (Fig. 7, Bioproject PRJNA528435; Supplementary Table 5d). Sesn1^-/-^ mice were used to show that loss of Sestrin1 aggravates disuse-induced muscle atrophy when compared with 6J mice (Segales et al., 2020); however, Sesn1^-/-^ mice were all *Nnt+* (Fig. 7, Bioproject PRJNA563889; Supplementary Table 5d).

**Fig. 7.**
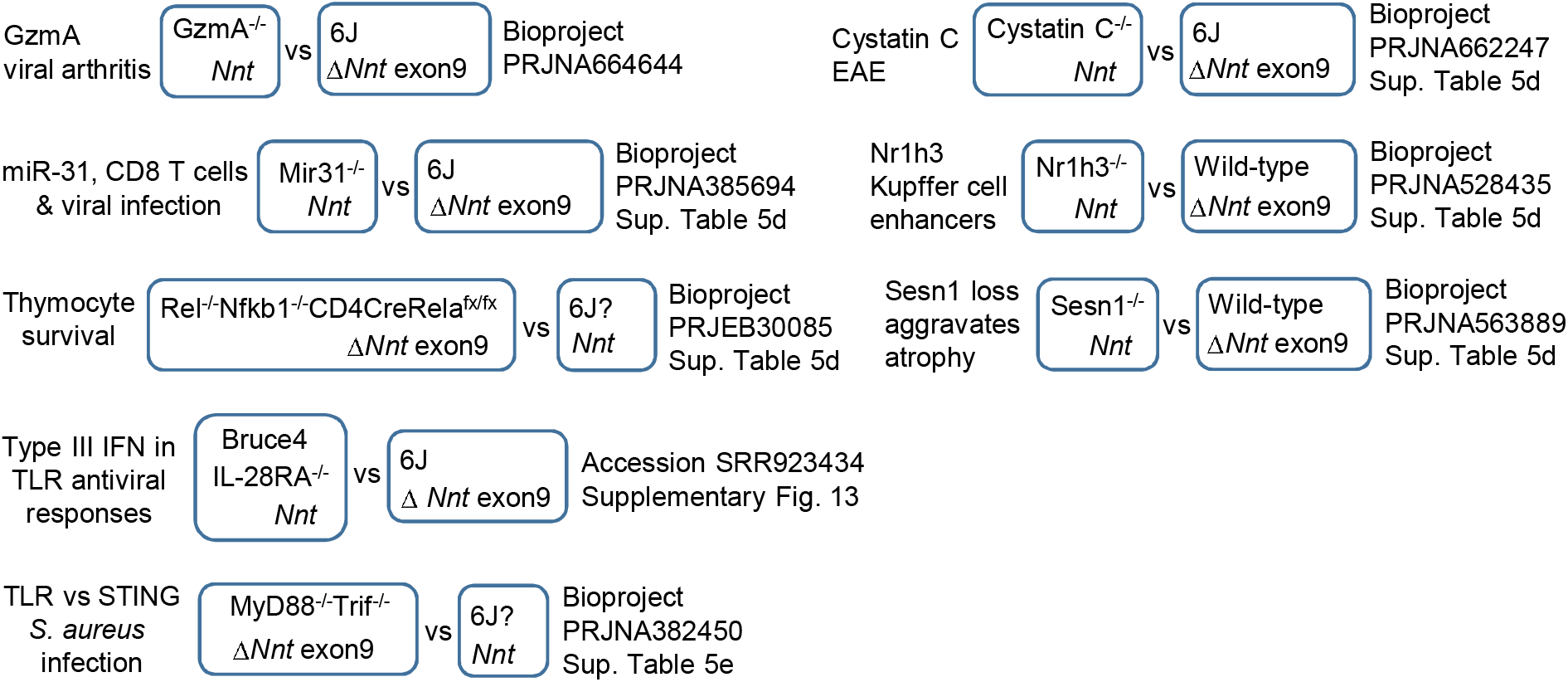
K-mer mining of Bioprojects where *Nnt^+/+^* mice were compared with *Nnt^-/-^* mice. The NCBI SRA data base was interrogated by k-mer mining for Bioprojects where some Accessions (listing 6J as the mouse strain) had reads compatible with a 6J background (reads for *Nnt* exon 2, but not exon 9) and other Accessions in that Bioproject (listing 6J as the mouse strain) had reads not compatible with a 6J background (reads for *Nnt* exon 2 and exon 9). The methodology is described in Supplementary Fig. 12a, validated by BLAST alignments (Supplementary Fig. 12b), with raw data in Supplementary Table 5. Selected projects are shown where publications and/or GEO xx

Of the 57 Bioprojects containing *Nnt+* and *Nnt+* Run Accessions, several contained comparisons in which one group contained a combination of *Nnt+* and *Nnt^-^* Run Accessions, while the other group(s) contained either all *Nnt+* or all *Nnt^-^* Run Accessions (Supplementary Table 5f).

Whether the *Nnt* differences (or other background gene differences) would have significantly affected the interpretation of the aforementioned studies remains to be established. Nevertheless, the data argues that differences in *Nnt* are widely under-appreciated in a range of research fields and have the potential to compromise a wide range of studies.

## Discussion

Despite a large body of literature (see Introduction) no clear consensus has emerged regarding the physiological function of GzmA. This lack of consensus might now, at least in part, be explained by the extensive use of GzmA^-/-^ mice (Supplemental Table 1). We show herein that this mouse strain is on a mixed 6J/6N background and contains full length *Nnt*, with the NNT protein involved in ROS regulation. For example, T cell activities can be regulated by ROS in a range of settings (Chen et al., 2016; Franchina et al., 2018; Murphy and Siegel, 2013; Yarosz and Chang, 2018), perhaps contributing to the controversy (Ebnet et al., 1995; Joeckel and Bird, 2014; Regner et al., 2009; Regner et al., 2011; Smyth et al., 2003) regarding the role of lymphocyte-derived GzmA as a mediator of cellular cytotoxicity (Pardo et al., 2002; Pardo et al., 2004; Shresta et al., 1999; Susanto et al., 2013). Whether all the pheontypes reported for GzmA^-/-^ mice were compromised by *Nnt* (or mixed the background) remains unclear and may require new experiments to resolve, similar to those described herein for CHIKV arthritis.

The data from GzmA^S211A^ mice (that were generated using CRISPR on a 6J background) represents the first *in vivo* assessment of the physiological function of GzmA without the confounding influence of differences in *Nnt* or other genes associated with the mixed genetic background. The results from this analysis support the view that the physiological activity of GzmA is mediated by its protease activity (Fig. 6b) (Schanoski et al., 2019). This is an important point because protease-independent functions have been documented for several proteases (Calhan and Seyrantepe, 2017; McNutt et al., 2007; Pu et al., 2019; Szabo et al., 2016). One of the proposed activities for GzmA, the binding to TLR9 and potentiation of TLR9 signaling (Shimizu et al., 2019) was not reported to require GzmA’s protease activity. However, using mice defective in TLR9 signaling, we were unable to find evidence that TLR9 is required for GzmA’s proinflammatory activity (Supplementary Fig. 14). Our studies also support the view that secreted extracellular GzmA has biological activity in the absence of perforin (or GzmB). This contrasts with the traditional view of GzmA as a mediator of cell death, which requires perforin to deliver GzmA to the cytoplasm, where a series of target molecules are cleaved (Liesche et al., 2018; Wu et al., 2019; Zhou et al., 2020). Cleavage of SET complex proteins (Mandrup-Poulsen, 2017; Mollah et al., 2017) would similarly require delivery of GzmA to the cytoplasm. Although we cannot formally exclude translocation of circulating GzmA into the cytoplasm via some unknown mechanism, the GzmA^S211A^ RNA-Seq experiment does ostensibly exclude perforin, as NK cells only produce perforin (and GzmB) protein after ≈24 hrs of appropriate stimulation (Fehniger et al., 2007).

Extracellular target candidates for GzmA include protease activated receptors (Hansen et al., 2005; Sower et al., 1996; Suidan et al., 1994; Suidan et al., 1996), and may also include pro-IL-1 (Hildebrand et al., 2014) (Fig. 6c), as pro-IL-1 can become extracellular when cells lyse (Afonina et al., 2015). Overall one might speculate that in such settings NK-derived GzmA synergizes with type I IFN responses to act as a systemic alarmin (Fig. 6b). GzmA^S211A^ mice will provide an invaluable tool for future studies into further refining our understanding of the role and molecular targets of GzmA.

The ability to reduce CHIKV arthritis in 6J mice with MitoTEMPO might suggest such antioxidant drugs have potential utility as anti-inflammatory treatments for alphaviral arthritides (Suhrbier et al., 2012; Zaid et al., 2020). However, MitoTEMPO treatment may simply be correcting the NNT defect in 6J mice by scavenging excess ROS arising from the loss of functional NNT (Ward et al., 2020). The argument that similar treatments would be effective in human diseases may thus not be compelling, given that most humans have a functional *Nnt* gene. Perhaps noteworthy is that many pre-clinical studies showing effective disease amelioration with MitoTEMPO treatments were conducted in 6J mice (Aoyama et al., 2012; Li et al., 2018; To et al., 2020; Vincent et al., 2020; Wu et al., 2020), whereas anti-oxidants have not shown clear benefits in human clinical trials (Casas et al., 2020; Jiang et al., 2020; Kovacic, 2020; Steinhubl, 2008).

The Jackson Laboratory generated the 6J inbred mouse strain in the 1920-1930s, with this mouse strain the most frequently used mouse strain in biomedical research. Although differences in the *Nnt* gene (or other background genes) have previously been reported as under-appreciated in metabolism research (Fontaine and Davis, 2016), the data herein and elsewhere (Bourdi et al., 2011; Ripoll et al., 2012; Vozenilek et al., 2018) argue that this issue extends to other areas of research, with ROS involved in many cellular processes (Gambhir et al., 2019; Sun et al., 2020). Of concern was that at least 26% of SRA Run Accessions and 38% of Bioprojects listing C57BL/6J as the mouse strain, had sequence data not consistent with a 6J background. Mouse strain listing errors or inadequate backcrossing to 6J would thus appear to be common for SRA RNA-Seq submissions. The full extent to which the NNT issue has complicated interpretation of knock-out mouse experiments remains to be addressed, but may require extensive new experiments such as those described herein for GzmA.

## Methods

### Ethics Statement and PC3/BSL3 certifications

All mouse work was conducted in accordance with the “Australian code for the care and use of animals for scientific purposes” as defined by the National Health and Medical Research Council of Australia. Mouse work was approved by the QIMR Berghofer Medical Research Institute animal ethics committee (P2235 A1606-618M), with infectious CHIKV work conducted in a biosafety level-3 (PC3) facility at the QIMR Berghofer MRI (Australian Department of Agriculture, Water and the Environment certification Q2326 and Office of the Gene Technology Regulator certification 3445).

### Cell lines and CHIKV

Vero (ATCC#: CCL-81) and C6/36 cells (ATCC# CRL-1660) were cultured as described (Nguyen et al., 2020). Cells were checked for mycoplasma using MycoAlert Mycoplasma Detection Kit (Lonza, Basel, Switzerland). FBS was checked for endotoxin contamination using RAW264.7-HIV-LTR-luc cells (Johnson et al., 2005) before purchase. CHIKV (isolate LR2006-OPY1; GenBank KT449801.1; DQ443544.2) was a kind gift from Dr. P. Roques (CEA, Fontenay-aux-Roses, France), was propagated in C6/36 cells, and titers determined by CCID_50_ assays (Nguyen et al., 2020). Virus was also checked for mycoplasma (La Linn et al., 1995).

### Mice and CHIKV infections

C57BL/6J mice were purchased from Animal Resources Centre (Canning Vale, WA, Australia). C57BL/6N mice were purchased from The Jackson Laboratory (Stock No. 005304). GzmA^-/-^ mice were generated as described (Ebnet et al., 1995) and were provided by the Peter MacCallum Cancer Centre, Melbourne, Victoria, Australia. GzmB^-/-^ mice were generated as described (Mullbacher et al., 1999) and were backcrossed onto C57BL/6J mice a total of 12 times and were provided by the Peter MacCallum Cancer Centre. The Australian Phenomics Network, Monash University, Melbourne, Australia used CRISPR to generate (i) GzmA^S211A^ mice on a 6J background and (ii) 6N^Δ*Nnt*8-12^ mice on a 6N background. IFNAR^-/-^ mice (Yan et al., 2020) were kindly provided by Dr P. Hertzog (Monash University, Melbourne, Australia). IL-28RA^-/-^ mice (Ank et al., 2008) were kindly provided by Bristol-Myers Squibb (Souza-Fonseca-Guimaraes et al., 2015). All GMO mice were bred in-house at QIMR Berghofer MRI.

Female mice 6-8 weeks old were infected with 10^4^ CCID_50_ CHIKV (isolate LR2006 OPY1) s.c. into each hind foot, with foot measurements and viremia determined as described (Gardner et al., 2010; Nguyen et al., 2020; Prow et al., 2018).

### RNA-Seq of feet CHIKV infection in GzmA^-/-^ versus CHIKV infection 6J mice

GzmA^-/-^ and 6J mice were infected with CHIKV, feet collected on day 6 and RNA samples prepared as described previously (Wilson et al., 2017). Library preparation and sequencing were conducted by the Australian Genome Research Facility (Melbourne, Australia) as described (Hazlewood et al., 2021). Briefly, cDNA libraries were prepared using a TruSeq RNA Sample Prep Kit (v2) (Illumina Inc. San Diego, USA), which included isolation of poly-adenylated RNA using oligo-dt beads. Paired end reads (100 bp) were generated using the Illumina HiSeq 2000 Sequencer (Illumina Inc.). Raw sequencing reads were assessed using FastQC (v0.11.8) (Simons, 2010) and MultiQC (v1.7) (Ewels et al., 2016) and trimmed using Cutadapt (v2.3) (Martin, 2011) to remove adapter sequences and low-quality bases. Trimmed reads were aligned to the GRCm38 primary assembly reference and the GENCODE M23 gene model using STAR aligner (v2.7.1a) (Dobin et al., 2012), with more than 95% of reads mapping to protein coding regions. Counts per gene were generated using RSEM (v1.3.1) (Li and Dewey, 2011) and differential expression analysis was undertaken using EdgeR in Galaxy (Varet et al., 2016) with default settings and no filters applied. Counts were normalized using the TMM method and modelled using the likelihood ratio test, glmLRT().

### RNA-Seq of poly(I:C) injection for GzmA^S211A^ versus 6J mice

Age matched female GzmA^S211A^ and 6J mice were injected i.v. with 250 μg of poly(I:C) in 150 μl of PBS. After 6 hours mice were euthanized, spleen and whole feet were harvested and RNA isolated as described previously (Prow et al., 2019; Wilson et al., 2017). Three RNA pools were generated for each mouse strain, whereby each pool contained equal amounts of RNA from 4 feet from 4 mice, or 2 spleens from 2 mice. RNA-Seq of polyadenylated RNA was then undertaken by in-house at QIMR Berghofer MRI. RNA integrity was assessed using the TapeStation system (Agilent Technologies) and libraries were prepared using the TruSeq Stranded mRNA library preparation kit (Illumina). Sequencing was performed on the Illumina Nextseq 550 platform with 75 bp paired end reads. Per base sequence quality for >92% bases was above Q30 for all samples. Raw sequencing reads were then processed as above.

### Bioinformatic analyses

Pathway analysis of differentially expressed genes in direct and indirect or direct only interactions were investigated using Ingenuity Pathway Analysis (IPA) (QIAGEN) (Shannon et al., 2003). Enrichment for biological processes, molecular functions, KEGG pathways and other gene ontology categories in DEG lists was elucidated using the STRING database (Szklarczyk et al., 2019) in Cytoscape (v3.7.2) (Shannon et al., 2003). Gene Set Enrichment Analysis (GSEA) (Subramanian et al., 2005) was performed on a desktop application (GSEA v4.0.3) (http://www.broadinstitute.org/gsea/) to look for enrichment of DEGs in full gene sets pre-ranked by fold change.

### Whole genome sequencing of GzmA^-/-^ mice

QIAamp DNA Micro Kit (QIAGEN) was used to purify genomic DNA from GzmA^-/-^ mice spleen as per manufacturer instructions. DNA was sent to Australian Genome Research Facility (AGRF) for Whole Genome Sequencing (WGS) using the Illumina NovaSeq platform with 150 bp paired end reads. The primary sequence data was generated using the Illumina bcl2fastq 2.20.0.422 pipeline. Reads were mapped to the mm10 genome assembly (GRCm38) using BWA-mem, and .bam files were provided. Mapped reads were viewed in Integrative Genomics Viewer (IGV) (Robinson et al., 2011) and 6N features identified manually based on previous publications (Mekada et al., 2015; Simon et al., 2013).

### Alignment to mouse genomes

Fastq files were generated as described or were downloaded from the Sequence Read Archive (SRA) using Aspera. Reads where trimmed using cutadapt (Martin, 2011), mapped using STAR aligner (v2.7.1a) for RNA-Seq or minimap 2 (Li, 2018) for WGS data. IGV was used to visualize reads mapping to the Nnt gene after mapping to the GRCm38 primary assembly reference for the truncated version of the gene and to the mouse C3H_HeJ_v1 reference (GCA_001632575.1) to observe full-length Nnt.

### *Nnt* RT PCR

RT PCR was undertaken essentially as described using the primer set (F1 AACAGTGCAAGGAGGTGGAC, R1 GTGCCAAGGTAAGCCACAAT) (Huang et al., 2006). RNA was extracted from testes using TRIzol (Sigma-Aldrich) according to manufacturer’s instructions. cDNA was generated using iScript cDNA Synthesis Kit (Bio-Rad) and Q5 Hot Start High-Fidelity DNA Polymerase (NEB) was used for PCR.

### MitoTEMPO treatment

Mice were injected i.v. daily, on the indicated day post CHIKV infection, with 62.5 μg of MitoTEMPO (Sigma-Aldrich) in 150 μl of PBS.

### K-mer mining

An exact-match (31 mer) k-mer mining approach was used to identify RNA-Seq read-files (Accessions) with C57BL/6J mice listed as the mouse strain, but where *Nnt* reads were incompatible with 6J background. Metadata associated with the National Center for Biotechnology Information’s SRA was screened using the Google Cloud Platform’s BigQuery service with the Structured Query Language (SQL) command: SELECT m.bioproject, m.acc, m.sample_name, m.platform, m.mbytes, m.mbases FROM nih-sra-datastore.sra.metadata as m, UNNEST (m.attributes) as a WHERE m.organism = ‘Mus musculus’ and m.assay_type = ‘RNA-Seq’ and a.v = ‘C57BL/6J’. Technical replicates for the same sample were collapsed by taking only the first accession for each Biosample. Accessions were then filtered on the basis of their compressed size so that only those between 200 Mb and 1500 Mb were retained; we found that read-files of this size provided adequate sequencing depth to detect *Nnt* exon reads. Accessions were sorted according to Bioproject and used as input for a bioinformatics pipeline executed on the Google Cloud Platform, which allowed access to the ‘SRA in the cloud’ database. A copy of our Bash script to automate the pipeline is available at https://github.com/CameronBishop/k-mer_mining_SRA. Accession read-files were converted to fastq format using fasterq-dump (SRA tool kit). BBduk version 38.87 (Bushnell, 2020) was used with default parameters to screen each read for sequence homology to exon 2 and exon 9 of the *Nnt* gene. Reads sharing at least one 31-mer with either exon were counted as a ‘match’ for that exon. Fastq files with at least ten matches to exon 2 and zero matches to exon 9 were classed as a consistent with a 6J genotype (truncated *Nnt*), while fastq files with at least ten matches to each exon were classed as a not consistent with a 6J genotype. Results were curated using BigQuery to confirm that for each Accession’s metadata entry, the ‘strain_sam’ field (or equivalent) of the metadata ‘Attributes’ table was listed as C57BL/6J.

Bioprojects were identified where some read-files contained exon 2 reads and no exon 9 reads, whereas others contained both exon 2 and exon 9 reads. The literature and Gene Expression Omnibus submissions associated with these Bioprojects were then consulted to identify Bioprojects where mice with full length *Nnt* had been compared mice with truncated *Nnt*.

### Determination of GzmA levels in mouse serum samples

Mouse serum was collected in Microvette 500 Z gel tubes (Sarstedt) with GzmA levels determined using a GzmA ELISA kit (MyBioSource, San Diego, CA, USA, MBS704766) according to manufacturer’s instructions.

### Statistics

Statistical analysis of experimental data was performed using IBM SPSS Statistics for Windows, Version 19.0. The t-test was performed when the difference in variances was <4, skewness was >-2 and kurtosis was <2, otherwise the Kolmogorov-Smirnov test was used.

## Acknowledgements

We thank the following staff at QIMR Berghofer MRI their assistance; animal house staff, Dr I Anraku (BSL3 facility manager) and Dr R. Johnston (Bioinformatics). We thank Dr Dion Kaiserman (Monash University, Australia) for supplying recombinant GzmA.

## DATA AVAILABILITY

All data is provided in the manuscript and accompanying supplementary files. Raw sequencing data generated for this publication has been deposited in the NCBI SRA: RNA-Seq, CHIKV arthritis GzmA^-/-^ vs 6J (PRJNA664644); RNA-Seq, poly(I:C) GzmA^S211A^ vs 6J (PRJNA666748); and WGS sequencing of GzmA^-/-^ mice (PRJNA664888).

## ETHICS STATEMENT

Mouse work was conducted in accordance with the “Australian code for the care and use of animals for scientific purposes” as defined by the National Health and Medical Research Council of Australia. Mouse work was approved by the QIMR Berghofer Medical Research Institute animal ethics committee (P2235).

## AUTHOR CONTRIBUTIONS

DJR, TTL, KY and EN undertook the experiments. TD and CB undertook the bioinformatics analyses. PIB provided vital reagents. AS and PIB conceptualized the study and obtained funding. AS wrote the manuscript with input from all the authors.

## SUPPLEMENTARY MATERIAL

The Supplementary Material for this article comprises Supplementary Figures (pdf) and Supplementary Tables (xlsx and doc).

