## Supplementary figures for "An inconvenient association between granzyme A and Nicotinamide Nucleotide Transhydrogenase"

### Slide 1
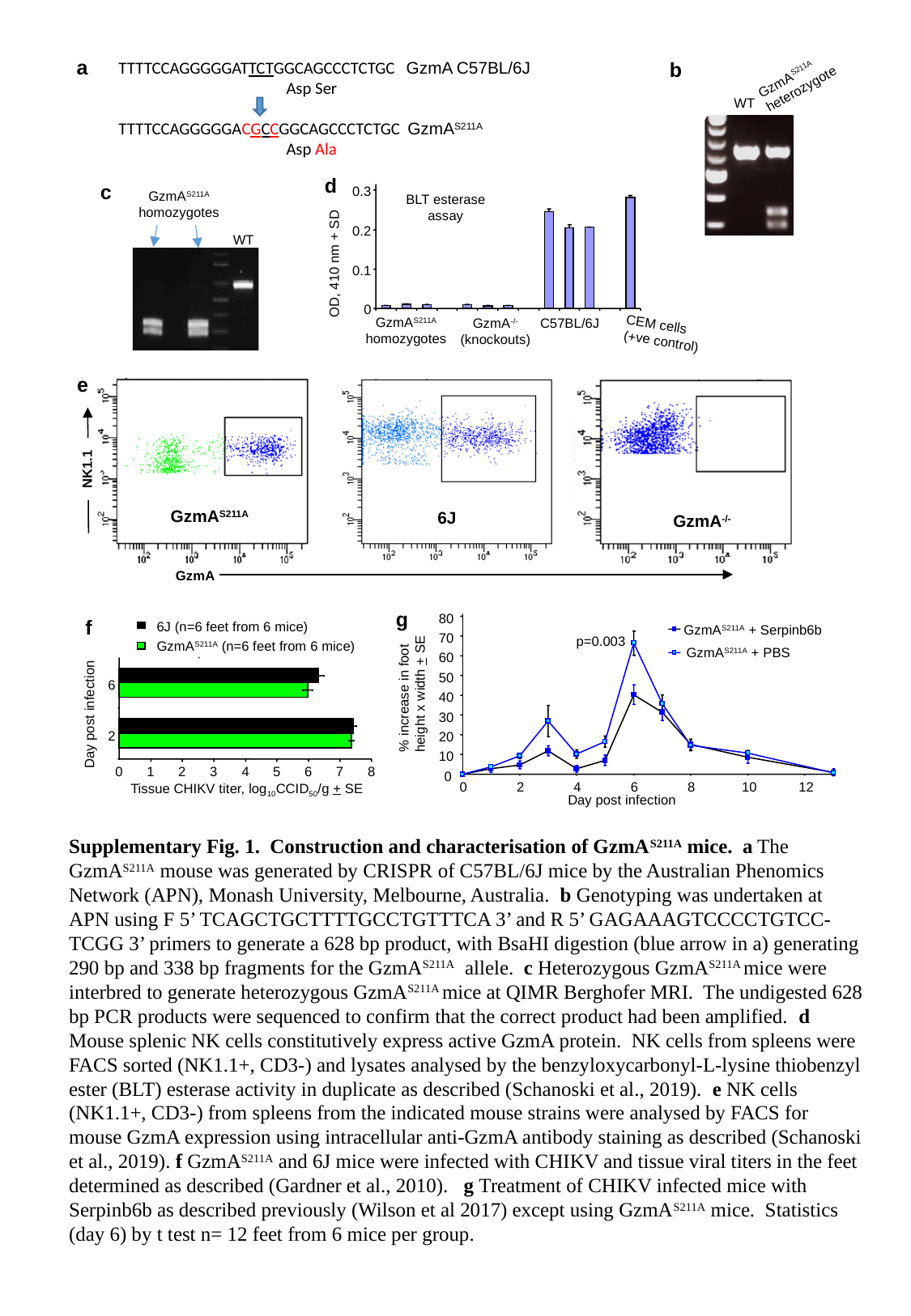

a
b
TTTTCCAGGGGGATTCTGGCAGCCCTCTGC GzmA C57BL/6J
	 Asp Ser
TTTTCCAGGGGGACGCCGGCAGCCCTCTGC GzmA^S211A^
	 Asp Ala
GzmA^S211A^
heterozygote
WT
d
c
GzmA^S211A^
homozygotes
0.3
BLT esterase
assay
0.2
OD, 410 nm + SD
0.1
0
GzmA^S211A^
homozygotes
C57BL/6J
GzmA^-/-^(knockouts)
CEM cells
(+ve control)
WT
e
NK1.1
GzmA^S211A^
6J
GzmA^-/-^
GzmA
g
f
6J (n=6 feet from 6 mice)
GzmA^S211A^ (n=6 feet from 6 mice)
6
Day post infection
2
0
1
2
3
4
5
6
7
8
Tissue CHIKV titer, log_10_CCID_50_/g + SE
80
70
60
50
40
30
20
10
0
GzmA^S211A^ + Serpinb6b
p=0.003
GzmA^S211A^ + PBS
% increase in foot
height x width + SE
0
2
4
6
8
10
12
Day post infection
Supplementary Fig. 1. Construction and characterisation of GzmA^S211A^ mice. a The GzmA^S211A^ mouse was generated by CRISPR of C57BL/6J mice by the Australian Phenomics Network (APN), Monash University, Melbourne, Australia. b Genotyping was undertaken at APN using F 5’ TCAGCTGCTTTTGCCTGTTTCA 3’ and R 5’ GAGAAAGTCCCCTGTCC-TCGG 3’ primers to generate a 628 bp product, with BsaHI digestion (blue arrow in a) generating 290 bp and 338 bp fragments for the GzmA^S211A^ allele. c Heterozygous GzmA^S211A^ mice were interbred to generate heterozygous GzmA^S211A^ mice at QIMR Berghofer MRI. The undigested 628 bp PCR products were sequenced to confirm that the correct product had been amplified. d Mouse splenic NK cells constitutively express active GzmA protein. NK cells from spleens were FACS sorted (NK1.1+, CD3-) and lysates analysed by the benzyloxycarbonyl-L-lysine thiobenzyl ester (BLT) esterase activity in duplicate as described (Schanoski et al., 2019). e NK cells (NK1.1+, CD3-) from spleens from the indicated mouse strains were analysed by FACS for mouse GzmA expression using intracellular anti-GzmA antibody staining as described (Schanoski et al., 2019). f GzmA^S211A^ and 6J mice were infected with CHIKV and tissue viral titers in the feet determined as described (Gardner et al., 2010). g Treatment of CHIKV infected mice with Serpinb6b as described previously (Wilson et al 2017) except using GzmA^S211A^ mice. Statistics (day 6) by t test n= 12 feet from 6 mice per group.

### Slide 2
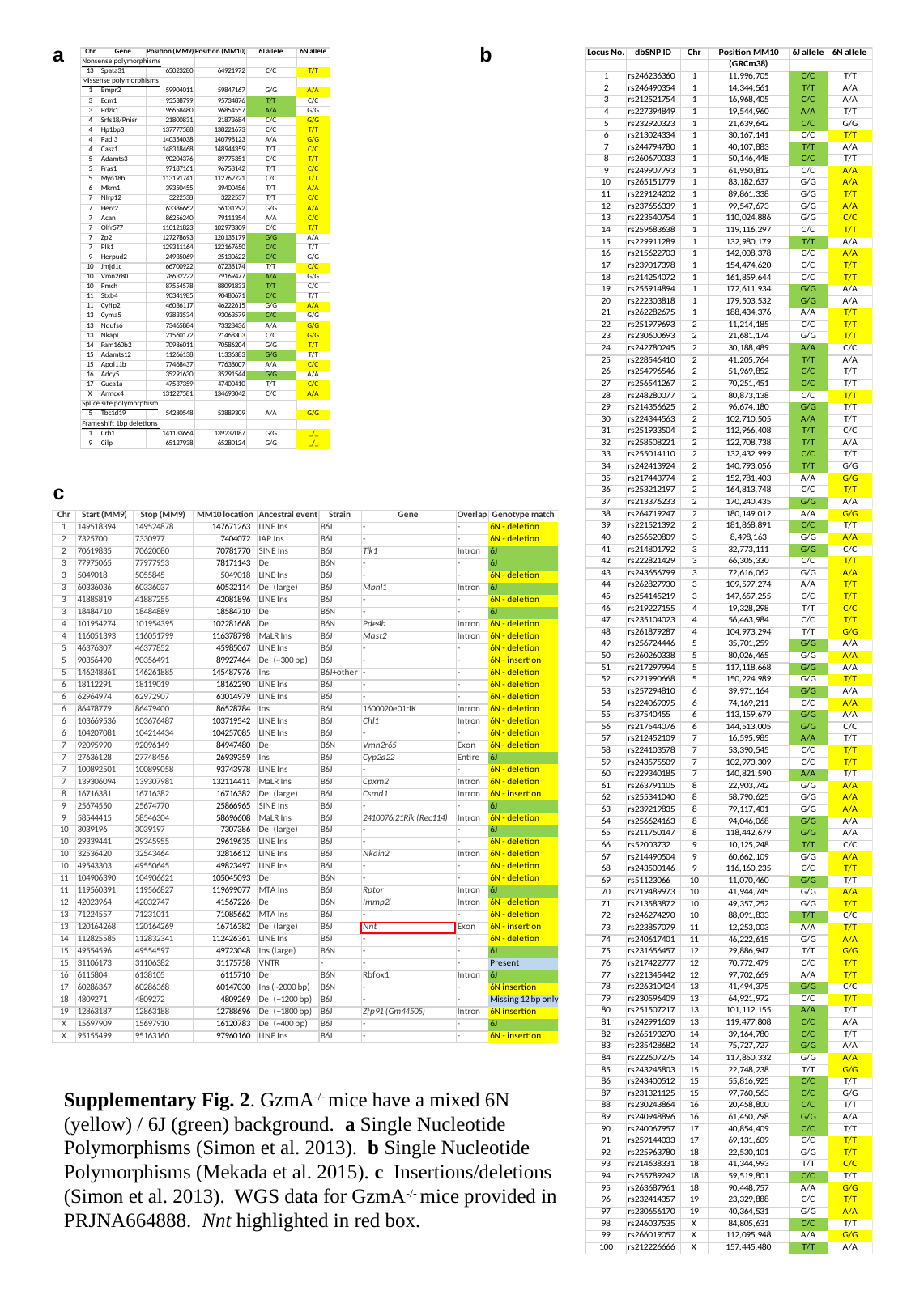

a
b
c
Supplementary Fig. 2. GzmA^-/-^ mice have a mixed 6N (yellow) / 6J (green) background. a Single Nucleotide Polymorphisms (Simon et al. 2013). b Single Nucleotide Polymorphisms (Mekada et al. 2015). c Insertions/deletions (Simon et al. 2013). WGS data for GzmA^-/-^ mice provided in PRJNA664888. Nnt highlighted in red box.

### Slide 3
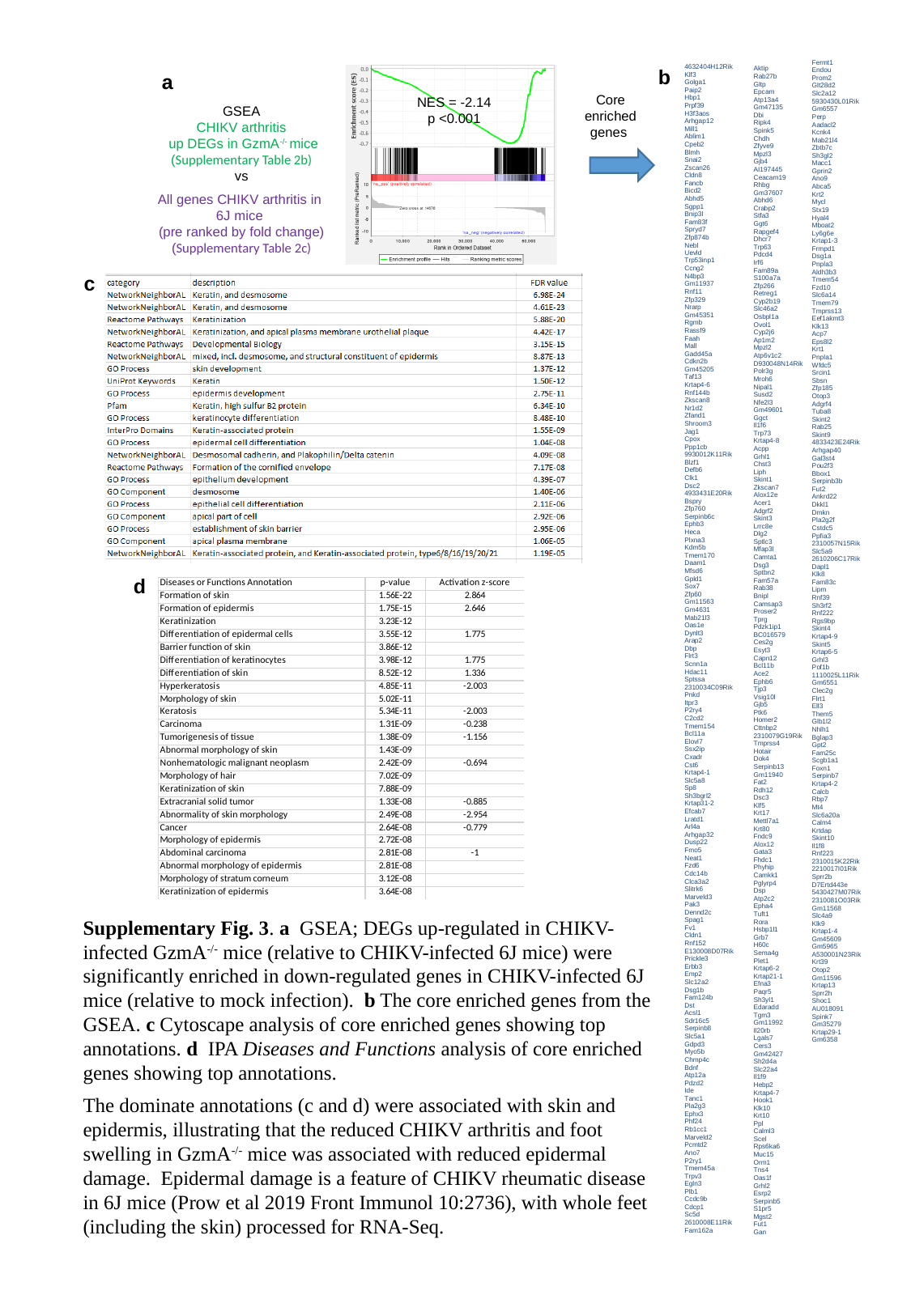

b
a
Core enriched genes
NES = -2.14
p <0.001
GSEA
CHIKV arthritis
 up DEGs in GzmA^-/-^ mice
(Supplementary Table 2b)
vs
All genes CHIKV arthritis in
6J mice
(pre ranked by fold change)
(Supplementary Table 2c)
c
d
Supplementary Fig. 3. a GSEA; DEGs up-regulated in CHIKV-infected GzmA^-/-^ mice (relative to CHIKV-infected 6J mice) were significantly enriched in down-regulated genes in CHIKV-infected 6J mice (relative to mock infection). b The core enriched genes from the GSEA. c Cytoscape analysis of core enriched genes showing top annotations. d IPA Diseases and Functions analysis of core enriched genes showing top annotations.
The dominate annotations (c and d) were associated with skin and epidermis, illustrating that the reduced CHIKV arthritis and foot swelling in GzmA^-/-^ mice was associated with reduced epidermal damage. Epidermal damage is a feature of CHIKV rheumatic disease in 6J mice (Prow et al 2019 Front Immunol 10:2736), with whole feet (including the skin) processed for RNA-Seq.

### Slide 4
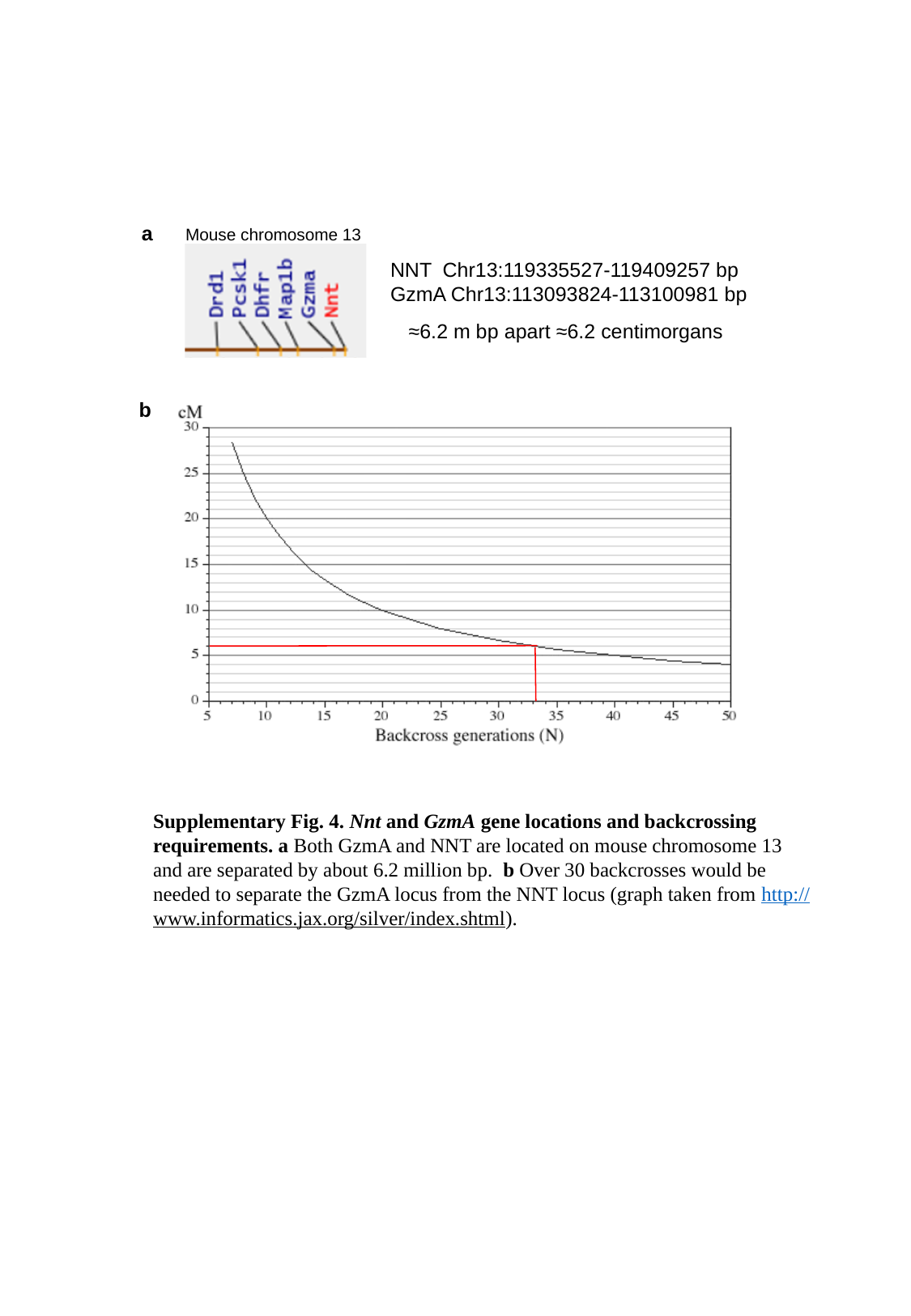

a
Mouse chromosome 13
NNT Chr13:119335527-119409257 bp
GzmA Chr13:113093824-113100981 bp
≈6.2 m bp apart ≈6.2 centimorgans
b
Supplementary Fig. 4. Nnt and GzmA gene locations and backcrossing requirements. a Both GzmA and NNT are located on mouse chromosome 13 and are separated by about 6.2 million bp. b Over 30 backcrosses would be needed to separate the GzmA locus from the NNT locus (graph taken from http://www.informatics.jax.org/silver/index.shtml).

### Slide 5
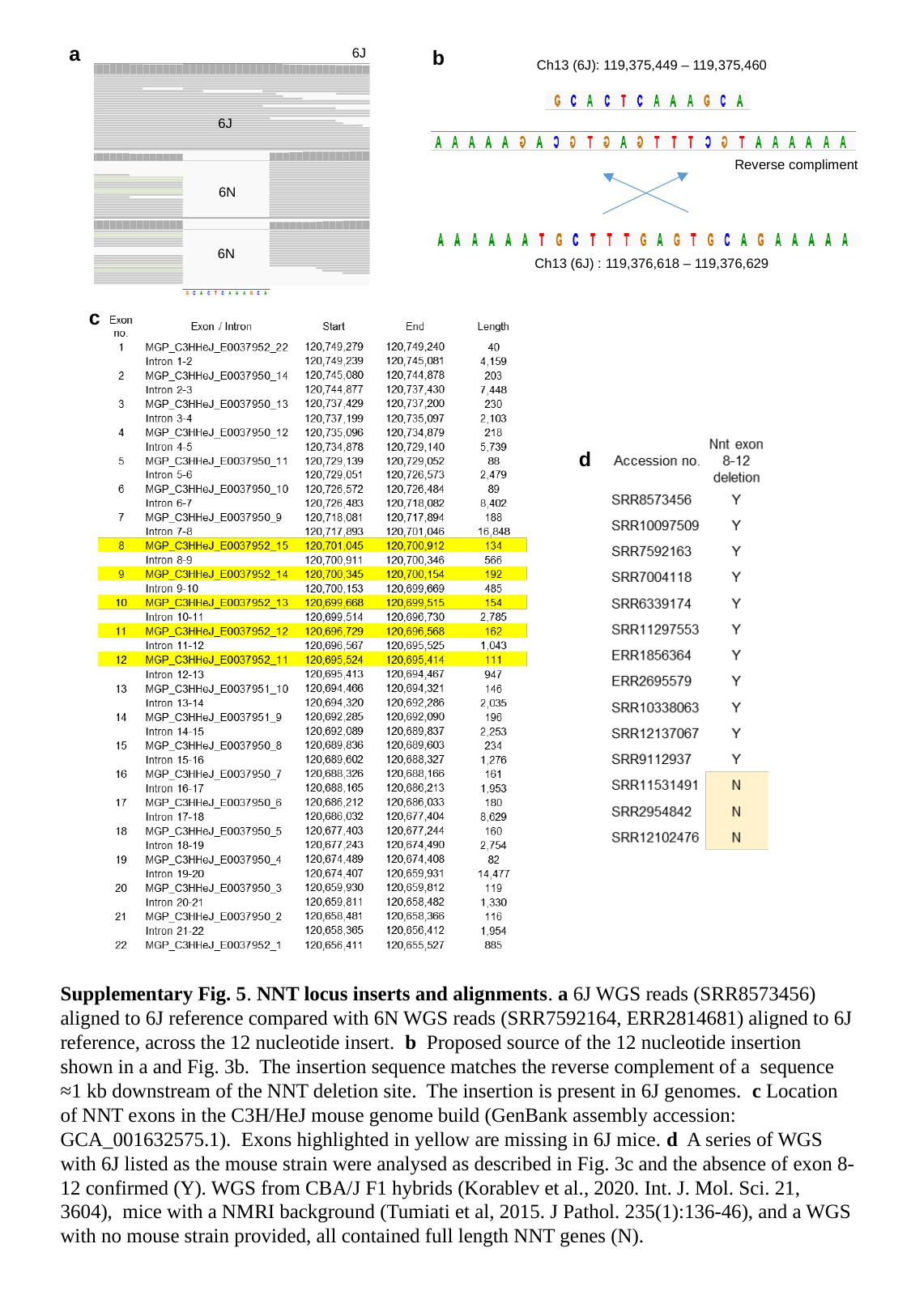

a
6J
b
Ch13 (6J): 119,375,449 – 119,375,460
6J
a
Reverse compliment
6N
6N
Ch13 (6J) : 119,376,618 – 119,376,629
c
d
Supplementary Fig. 5. NNT locus inserts and alignments. a 6J WGS reads (SRR8573456) aligned to 6J reference compared with 6N WGS reads (SRR7592164, ERR2814681) aligned to 6J reference, across the 12 nucleotide insert. b Proposed source of the 12 nucleotide insertion shown in a and Fig. 3b. The insertion sequence matches the reverse complement of a sequence ≈1 kb downstream of the NNT deletion site. The insertion is present in 6J genomes. c Location of NNT exons in the C3H/HeJ mouse genome build (GenBank assembly accession: GCA_001632575.1). Exons highlighted in yellow are missing in 6J mice. d A series of WGS with 6J listed as the mouse strain were analysed as described in Fig. 3c and the absence of exon 8-12 confirmed (Y). WGS from CBA/J F1 hybrids (Korablev et al., 2020. Int. J. Mol. Sci. 21, 3604), mice with a NMRI background (Tumiati et al, 2015. J Pathol. 235(1):136-46), and a WGS with no mouse strain provided, all contained full length NNT genes (N).

### Slide 6
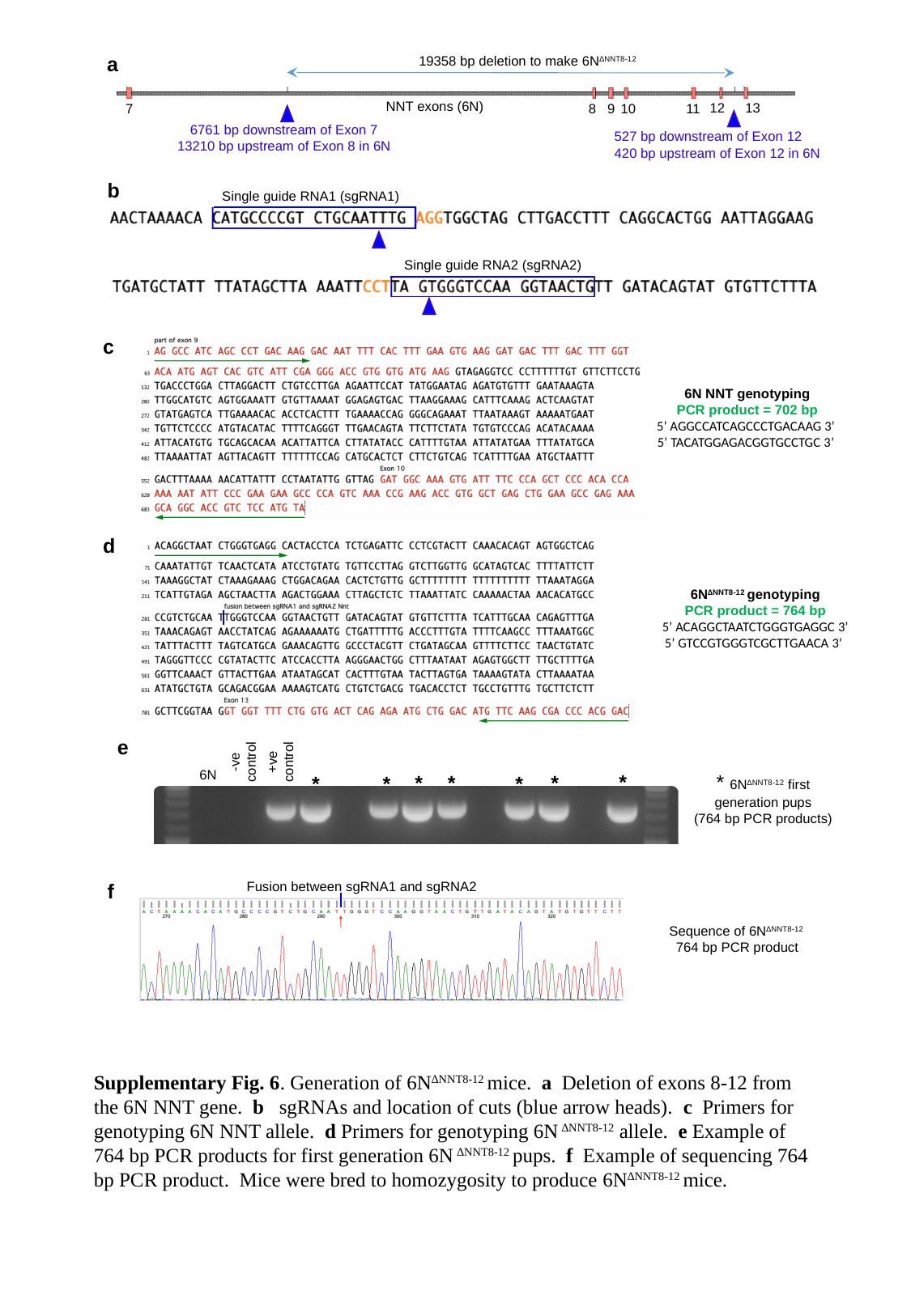

a
19358 bp deletion to make 6N^∆NNT8-12^
NNT exons (6N)
12
13
7
8
9
10
11
6761 bp downstream of Exon 7
13210 bp upstream of Exon 8 in 6N
527 bp downstream of Exon 12
420 bp upstream of Exon 12 in 6N
b
Single guide RNA1 (sgRNA1)
Single guide RNA2 (sgRNA2)
c
6N NNT genotyping
PCR product = 702 bp
5’ AGGCCATCAGCCCTGACAAG 3’
5’ TACATGGAGACGGTGCCTGC 3’
d
6N^∆NNT8-12^ genotyping
PCR product = 764 bp
5’ ACAGGCTAATCTGGGTGAGGC 3’
5’ GTCCGTGGGTCGCTTGAACA 3’
-ve
control
+ve
control
6N
*
* 6N^∆NNT8-12^ first generation pups
(764 bp PCR products)
*
*
*
*
*
*
e
Fusion between sgRNA1 and sgRNA2
f
Sequence of 6N^∆NNT8-12^
764 bp PCR product
Supplementary Fig. 6. Generation of 6N^∆NNT8-12^ mice. a Deletion of exons 8-12 from the 6N NNT gene. b sgRNAs and location of cuts (blue arrow heads). c Primers for genotyping 6N NNT allele. d Primers for genotyping 6N ^∆NNT8-12^ allele. e Example of 764 bp PCR products for first generation 6N ^∆NNT8-12^ pups. f Example of sequencing 764 bp PCR product. Mice were bred to homozygosity to produce 6N^∆NNT8-12^ mice.

### Slide 7
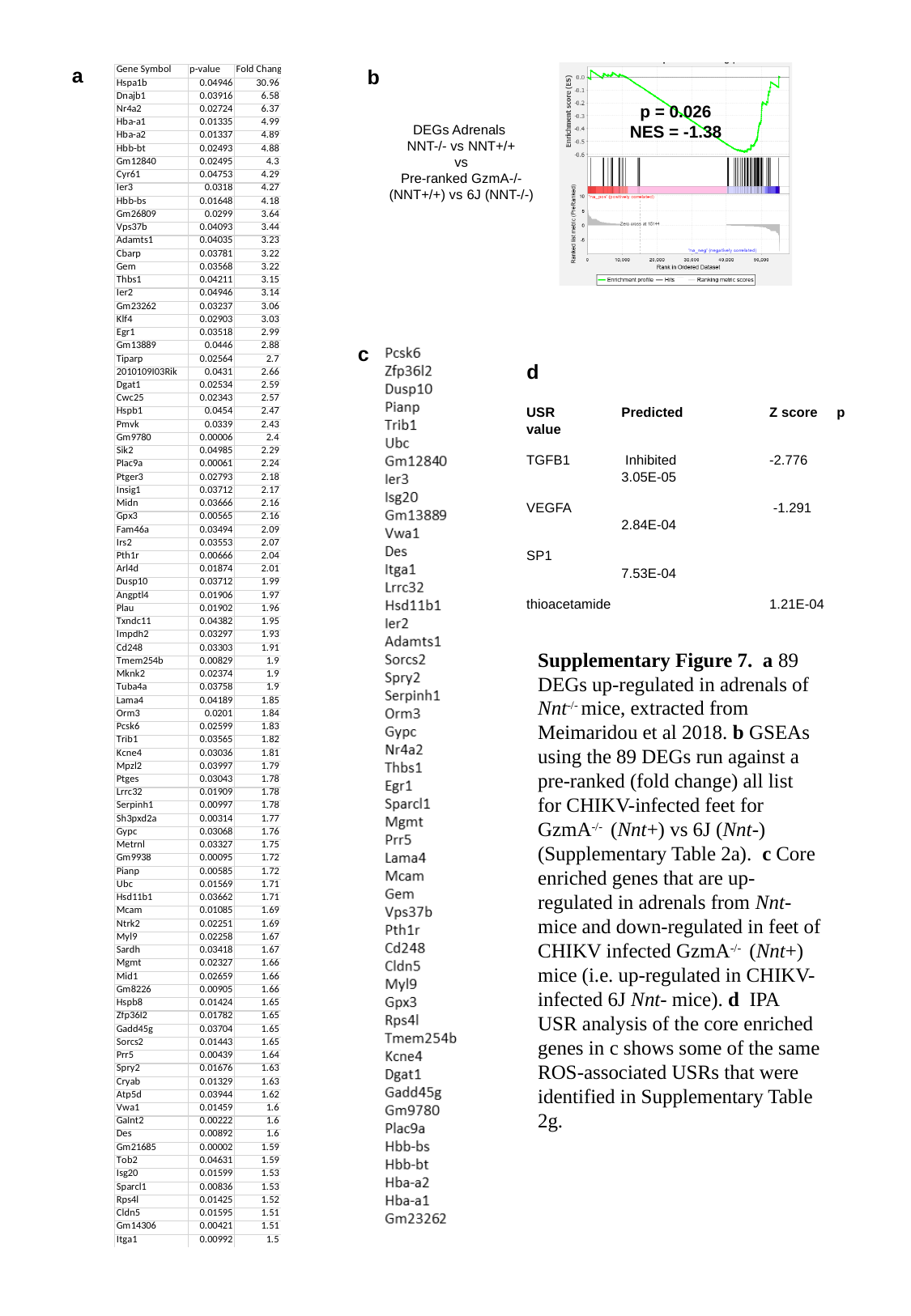

a
b
p = 0.026
NES = -1.38
DEGs Adrenals
NNT-/- vs NNT+/+
vs
Pre-ranked GzmA-/- (NNT+/+) vs 6J (NNT-/-)
c
d
USR	Predicted	Z score p value
TGFB1	 Inhibited	-2.776	3.05E-05
VEGFA	 		 -1.291	2.84E-04
SP1	 		 	7.53E-04
thioacetamide	 	 	1.21E-04
Supplementary Figure 7. a 89 DEGs up-regulated in adrenals of Nnt^-/-^ mice, extracted from Meimaridou et al 2018. b GSEAs using the 89 DEGs run against a pre-ranked (fold change) all list for CHIKV-infected feet for GzmA^-/-^ (Nnt+) vs 6J (Nnt-) (Supplementary Table 2a). c Core enriched genes that are up-regulated in adrenals from Nnt- mice and down-regulated in feet of CHIKV infected GzmA^-/-^ (Nnt+) mice (i.e. up-regulated in CHIKV-infected 6J Nnt- mice). d IPA USR analysis of the core enriched genes in c shows some of the same ROS-associated USRs that were identified in Supplementary Table 2g.

### Slide 8
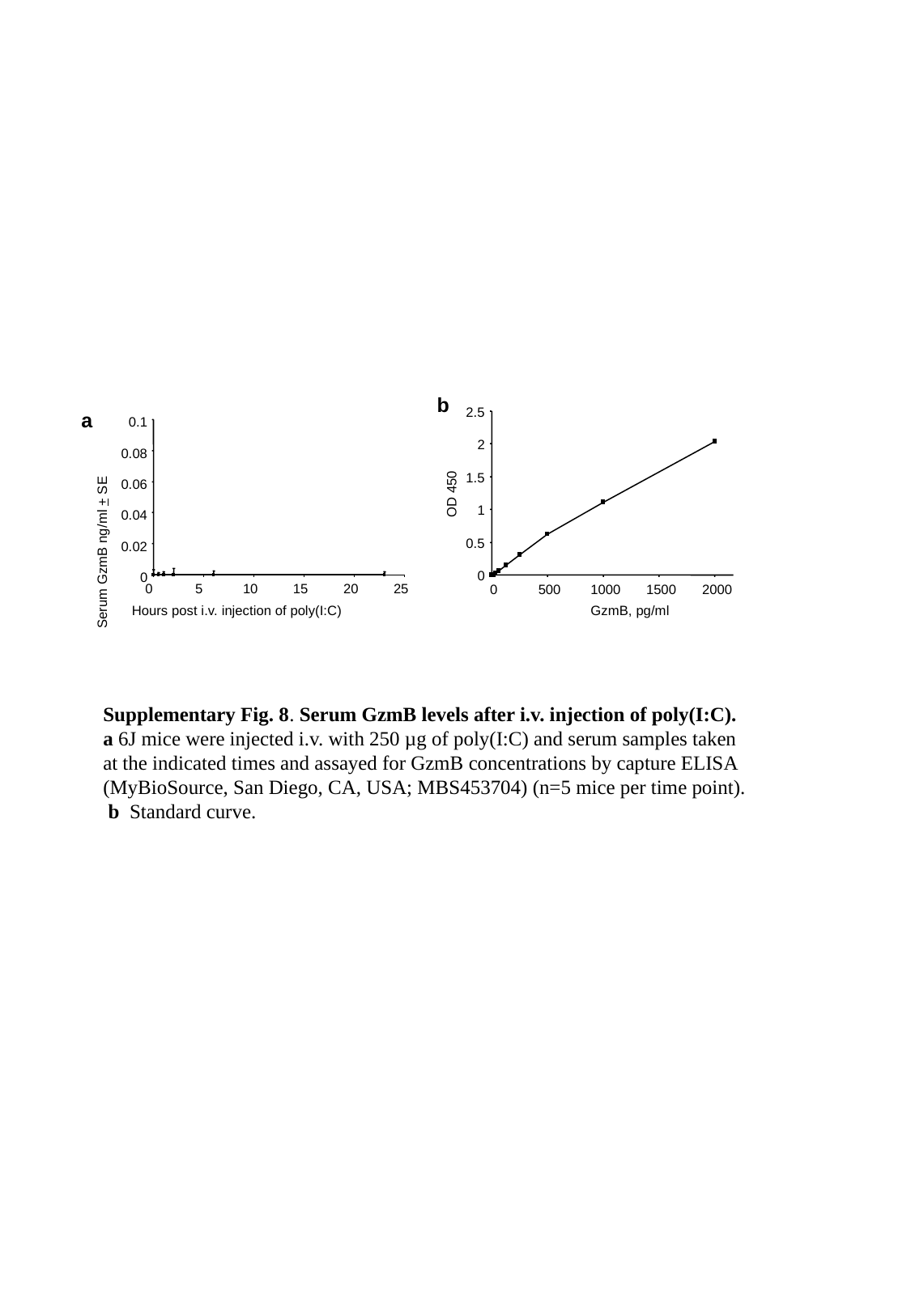

b
a
2.5
2
1.5
1
0.5
0
0.1
0.08
0.06
0.04
0.02
0
OD 450
Serum GzmB ng/ml + SE
0
5
10
15
20
25
0
500
1000
1500
2000
Hours post i.v. injection of poly(I:C)
GzmB, pg/ml
Supplementary Fig. 8. Serum GzmB levels after i.v. injection of poly(I:C). a 6J mice were injected i.v. with 250 µg of poly(I:C) and serum samples taken at the indicated times and assayed for GzmB concentrations by capture ELISA (MyBioSource, San Diego, CA, USA; MBS453704) (n=5 mice per time point). b Standard curve.

### Slide 9
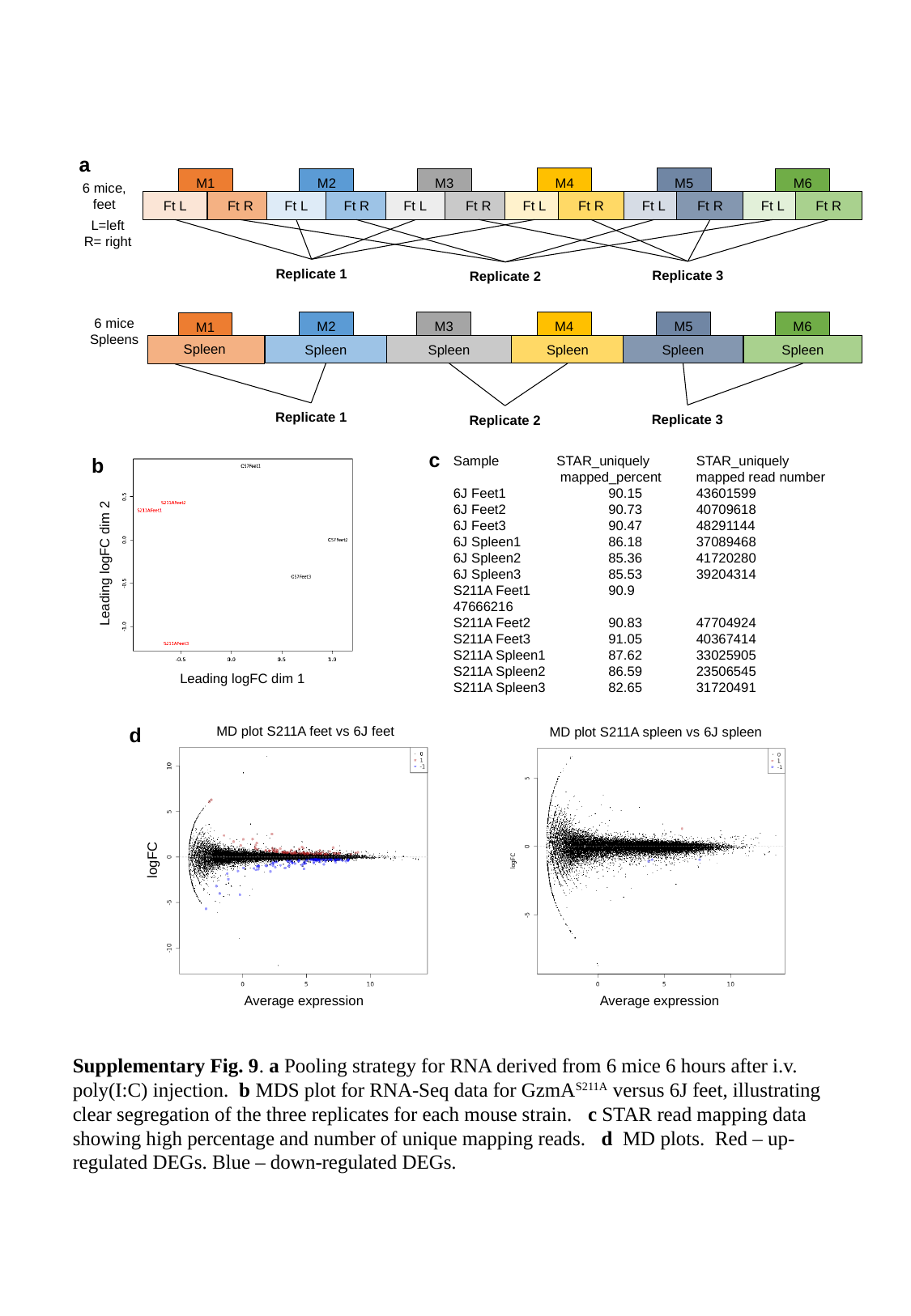

a
M5
M4
M2
M3
M6
M1
6 mice, feet
Ft R
Ft R
Ft L
Ft L
Ft L
Ft R
Ft L
Ft R
Ft R
Ft L
Ft R
Ft L
L=left
R= right
Replicate 1
Replicate 3
Replicate 2
6 mice
Spleens
M5
M4
M2
M3
M6
M1
Spleen
Spleen
Spleen
Spleen
Spleen
Spleen
Replicate 1
Replicate 3
Replicate 2
Leading logFC dim 2
Leading logFC dim 1
c
b
Sample STAR_uniquely	STAR_uniquely 	mapped_percent mapped read number
6J Feet1	90.15	43601599
6J Feet2	90.73	40709618
6J Feet3	90.47	48291144
6J Spleen1	86.18	37089468
6J Spleen2	85.36	41720280
6J Spleen3	85.53	39204314
S211A Feet1	90.9		47666216
S211A Feet2	90.83	47704924
S211A Feet3	91.05	40367414
S211A Spleen1	87.62	33025905
S211A Spleen2	86.59	23506545
S211A Spleen3	82.65	31720491
d
MD plot S211A feet vs 6J feet
MD plot S211A spleen vs 6J spleen
logFC
Average expression
Average expression
Supplementary Fig. 9. a Pooling strategy for RNA derived from 6 mice 6 hours after i.v. poly(I:C) injection. b MDS plot for RNA-Seq data for GzmA^S211A^ versus 6J feet, illustrating clear segregation of the three replicates for each mouse strain. c STAR read mapping data showing high percentage and number of unique mapping reads. d MD plots. Red – up-regulated DEGs. Blue – down-regulated DEGs.

### Slide 10
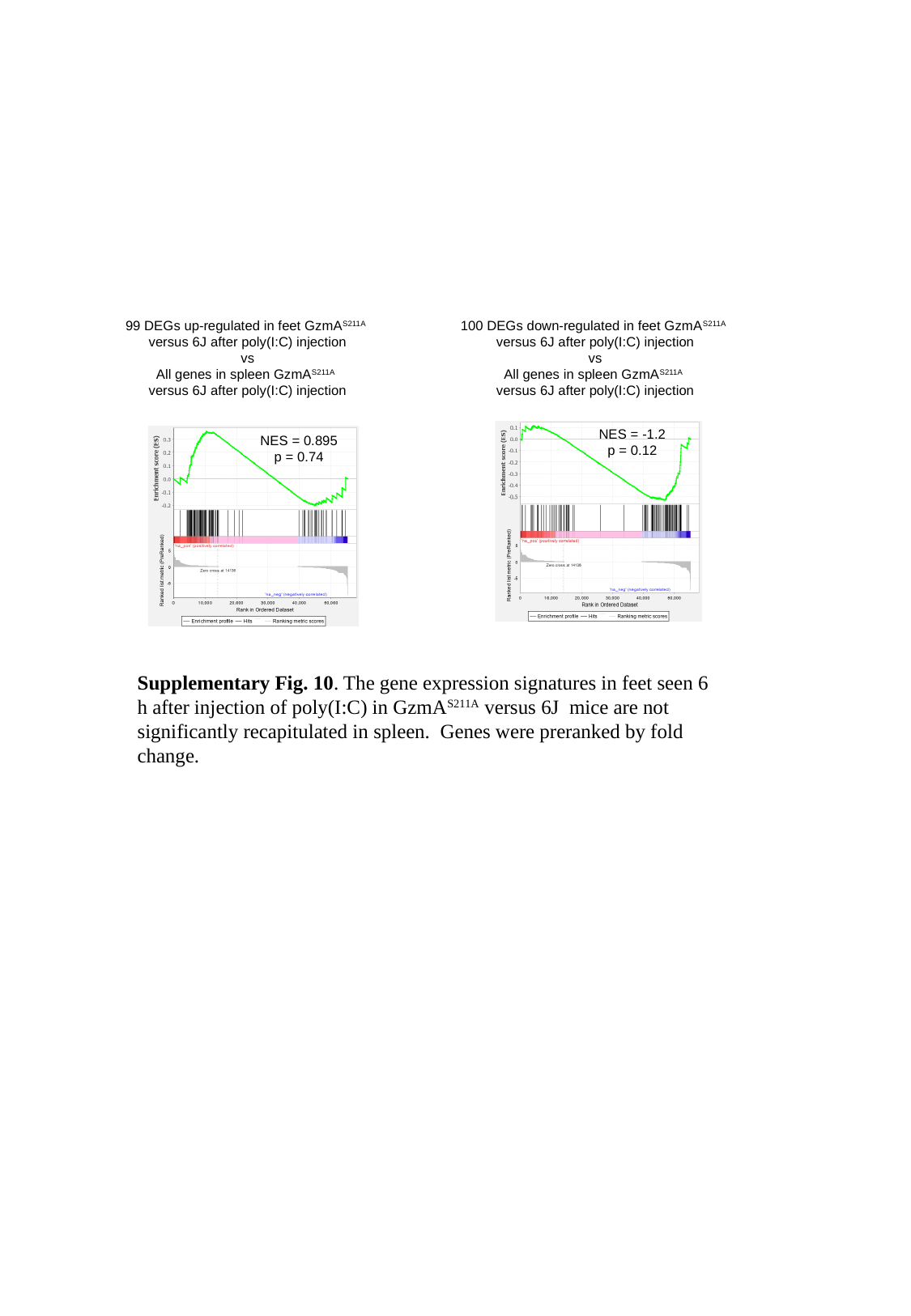

99 DEGs up-regulated in feet GzmA^S211A^
versus 6J after poly(I:C) injection
vs
All genes in spleen GzmA^S211A^
versus 6J after poly(I:C) injection
100 DEGs down-regulated in feet GzmA^S211A^
versus 6J after poly(I:C) injection
vs
All genes in spleen GzmA^S211A^
versus 6J after poly(I:C) injection
NES = -1.2
p = 0.12
NES = 0.895
p = 0.74
Supplementary Fig. 10. The gene expression signatures in feet seen 6 h after injection of poly(I:C) in GzmA^S211A^ versus 6J mice are not significantly recapitulated in spleen. Genes were preranked by fold change.

### Slide 11
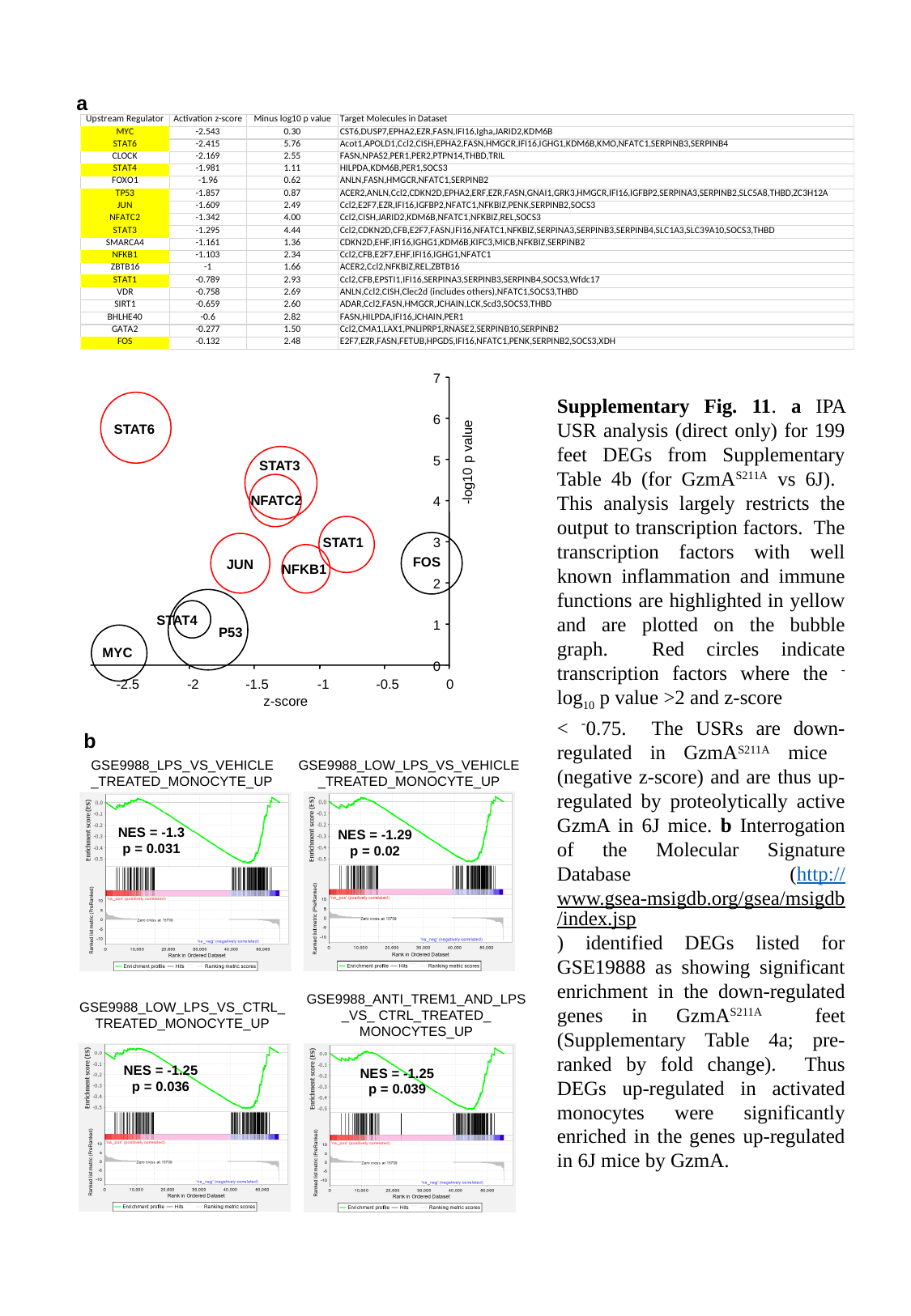

a
7
6
STAT6
-log10 p value
STAT3
5
NFATC2
4
STAT1
3
FOS
JUN
NFKB1
2
STAT4
1
P53
MYC
0
-2.5
-2
-1.5
-1
-0.5
0
z-score
Supplementary Fig. 11. a IPA USR analysis (direct only) for 199 feet DEGs from Supplementary Table 4b (for GzmA^S211A^ vs 6J). This analysis largely restricts the output to transcription factors. The transcription factors with well known inflammation and immune functions are highlighted in yellow and are plotted on the bubble graph. Red circles indicate transcription factors where the ^-^log_10_ p value >2 and z-score
< ^-^0.75. The USRs are down-regulated in GzmA^S211A^ mice (negative z-score) and are thus up-regulated by proteolytically active GzmA in 6J mice. b Interrogation of the Molecular Signature Database (http://www.gsea-msigdb.org/gsea/msigdb/index.jsp) identified DEGs listed for GSE19888 as showing significant enrichment in the down-regulated genes in GzmA^S211A^ feet (Supplementary Table 4a; pre-ranked by fold change). Thus DEGs up-regulated in activated monocytes were significantly enriched in the genes up-regulated in 6J mice by GzmA.
b
GSE9988_LPS_VS_VEHICLE
_TREATED_MONOCYTE_UP
GSE9988_LOW_LPS_VS_VEHICLE
_TREATED_MONOCYTE_UP
NES = -1.3
p = 0.031
NES = -1.29
p = 0.02
GSE9988_ANTI_TREM1_AND_LPS
_VS_ CTRL_TREATED_
MONOCYTES_UP
GSE9988_LOW_LPS_VS_CTRL_
TREATED_MONOCYTE_UP
NES = -1.25
p = 0.036
NES = -1.25
p = 0.039

### Slide 12
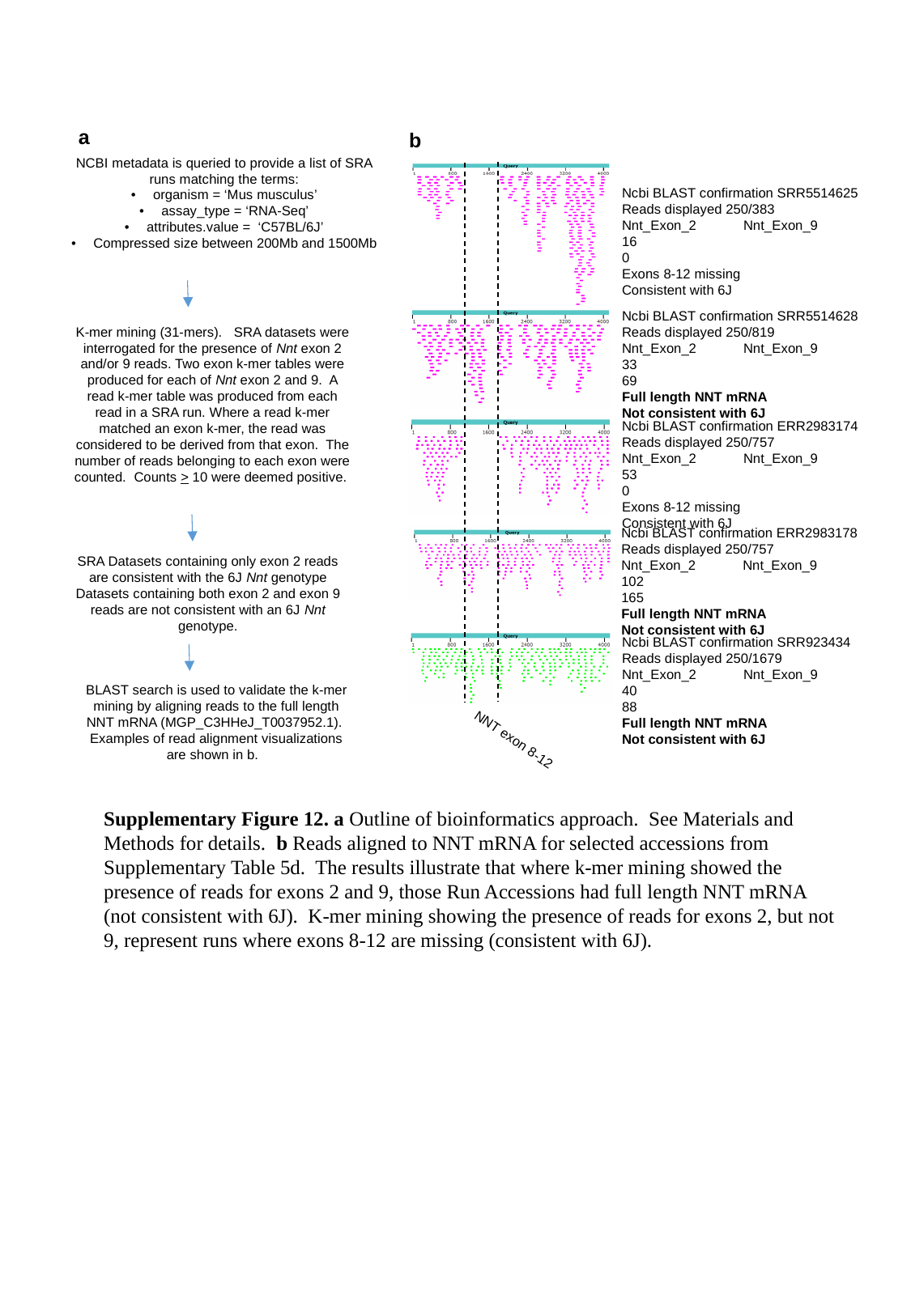

a
b
NCBI metadata is queried to provide a list of SRA runs matching the terms:
organism = ‘Mus musculus’
assay_type = ‘RNA-Seq’
attributes.value = ‘C57BL/6J’
Compressed size between 200Mb and 1500Mb
Ncbi BLAST confirmation SRR5514625
Reads displayed 250/383
Nnt_Exon_2	Nnt_Exon_9
16		0
Exons 8-12 missing
Consistent with 6J
Ncbi BLAST confirmation SRR5514628
Reads displayed 250/819
Nnt_Exon_2	Nnt_Exon_9
33		69
Full length NNT mRNA
Not consistent with 6J
K-mer mining (31-mers). SRA datasets were interrogated for the presence of Nnt exon 2 and/or 9 reads. Two exon k-mer tables were produced for each of Nnt exon 2 and 9. A read k-mer table was produced from each read in a SRA run. Where a read k-mer matched an exon k-mer, the read was considered to be derived from that exon. The number of reads belonging to each exon were counted. Counts > 10 were deemed positive.
Ncbi BLAST confirmation ERR2983174
Reads displayed 250/757
Nnt_Exon_2	Nnt_Exon_9
53		0
Exons 8-12 missing
Consistent with 6J
Ncbi BLAST confirmation ERR2983178
Reads displayed 250/757
Nnt_Exon_2	Nnt_Exon_9
102		165
Full length NNT mRNA
Not consistent with 6J
SRA Datasets containing only exon 2 reads are consistent with the 6J Nnt genotype
Datasets containing both exon 2 and exon 9 reads are not consistent with an 6J Nnt genotype.
Ncbi BLAST confirmation SRR923434
Reads displayed 250/1679
Nnt_Exon_2	Nnt_Exon_9
40		88
Full length NNT mRNA
Not consistent with 6J
BLAST search is used to validate the k-mer mining by aligning reads to the full length NNT mRNA (MGP_C3HHeJ_T0037952.1). Examples of read alignment visualizations are shown in b.
NNT exon 8-12
Supplementary Figure 12. a Outline of bioinformatics approach. See Materials and Methods for details. b Reads aligned to NNT mRNA for selected accessions from Supplementary Table 5d. The results illustrate that where k-mer mining showed the presence of reads for exons 2 and 9, those Run Accessions had full length NNT mRNA (not consistent with 6J). K-mer mining showing the presence of reads for exons 2, but not 9, represent runs where exons 8-12 are missing (consistent with 6J).

### Slide 13
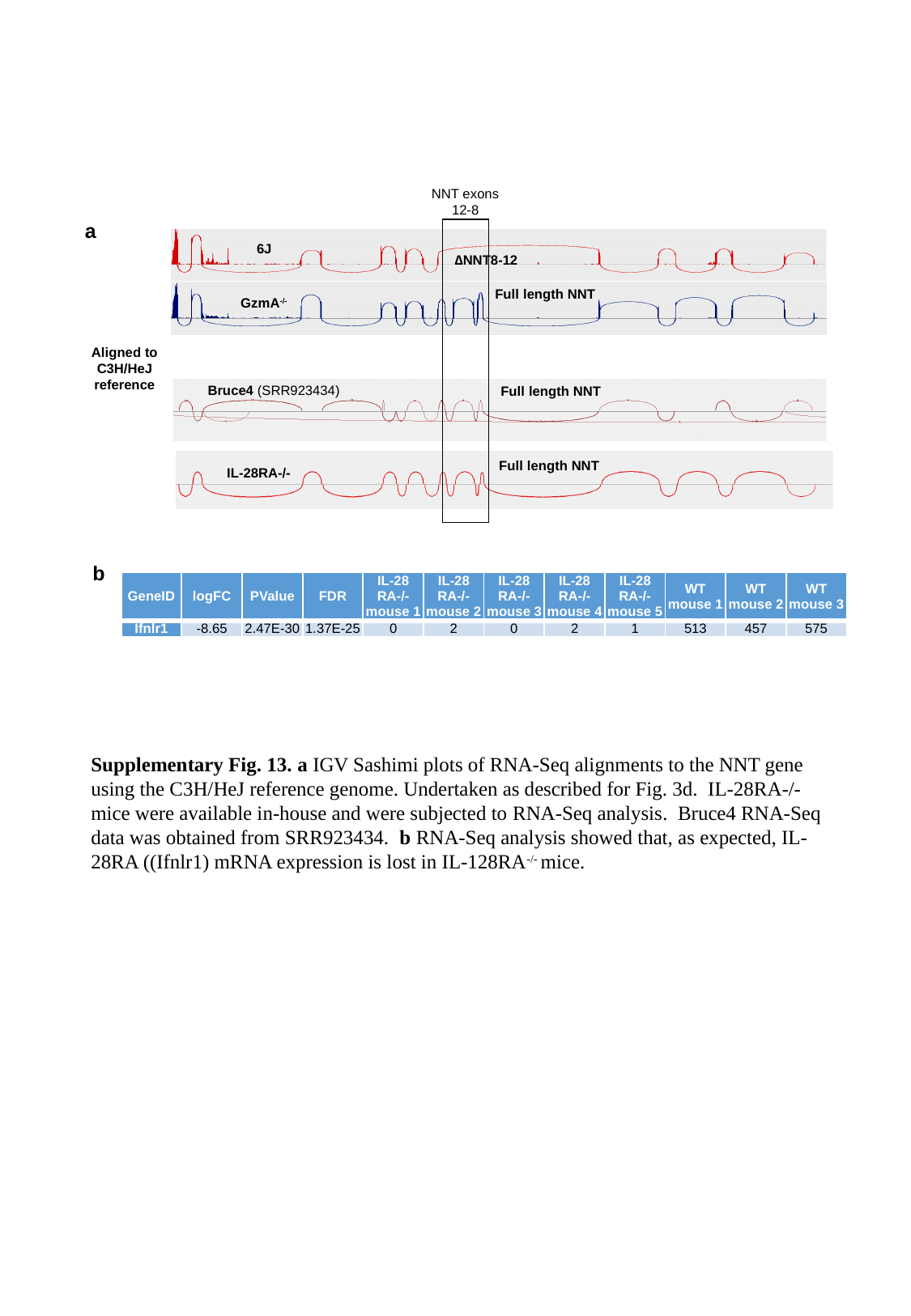

NNT exons
12-8
a
6J
∆NNT8-12
Full length NNT
GzmA^-/-^
Aligned to
C3H/HeJ
reference
Bruce4 (SRR923434)
Full length NNT
Full length NNT
IL-28RA-/-
b
| GeneID | logFC | PValue | FDR | IL-28 RA-/- mouse 1 | IL-28 RA-/- mouse 2 | IL-28 RA-/- mouse 3 | IL-28 RA-/- mouse 4 | IL-28 RA-/- mouse 5 | WT mouse 1 | WT mouse 2 | WT mouse 3 |
| --- | --- | --- | --- | --- | --- | --- | --- | --- | --- | --- | --- |
| Ifnlr1 | -8.65 | 2.47E-30 | 1.37E-25 | 0 | 2 | 0 | 2 | 1 | 513 | 457 | 575 |
Supplementary Fig. 13. a IGV Sashimi plots of RNA-Seq alignments to the NNT gene using the C3H/HeJ reference genome. Undertaken as described for Fig. 3d. IL-28RA-/- mice were available in-house and were subjected to RNA-Seq analysis. Bruce4 RNA-Seq data was obtained from SRR923434. b RNA-Seq analysis showed that, as expected, IL-28RA ((Ifnlr1) mRNA expression is lost in IL-128RA^-/-^ mice.

### Slide 14
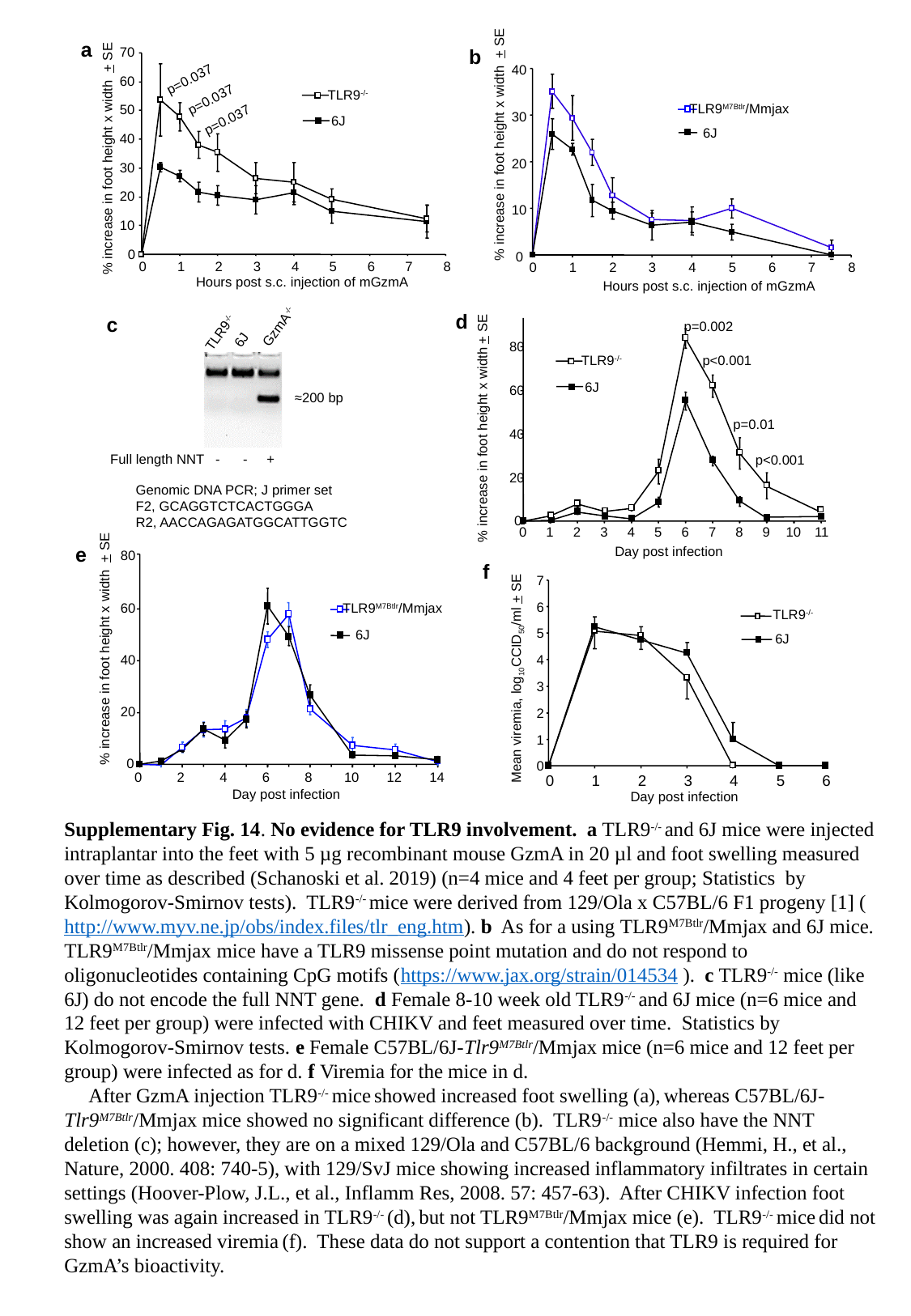

40
TLR9^M7Btlr^/Mmjax
6J
30
% increase in foot height x width + SE
20
10
0
0
1
2
3
4
5
6
7
8
Hours post s.c. injection of mGzmA
a
b
70
60
50
40
30
20
10
0
p=0.037
p=0.037
TLR9^-/-^
6J
p=0.037
% increase in foot height x width + SE
0
1
2
3
4
5
6
7
8
Hours post s.c. injection of mGzmA
c
GzmA^-/-^
TLR9^-/-^
6J
≈200 bp
Full length NNT - - +
Genomic DNA PCR; J primer set
F2, GCAGGTCTCACTGGGA
R2, AACCAGAGATGGCATTGGTC
d
p=0.002
80
60
40
20
0
p<0.001
TLR9^-/-^
6J
p=0.01
% increase in foot height x width + SE
p<0.001
0
1
2
3
4
5
6
7
8
9
10
11
e
Day post infection
80
60
40
20
0
f
7
6
5
4
3
2
1
0
TLR9^-/-^
6J
Mean viremia, log_10_CCID_50_/ml + SE
0
1
2
3
4
5
6
Day post infection
TLR9^M7Btlr^/Mmjax
6J
% increase in foot height x width + SE
0
2
4
6
8
10
12
14
Day post infection
Supplementary Fig. 14. No evidence for TLR9 involvement. a TLR9^-/-^ and 6J mice were injected intraplantar into the feet with 5 µg recombinant mouse GzmA in 20 µl and foot swelling measured over time as described (Schanoski et al. 2019) (n=4 mice and 4 feet per group; Statistics by Kolmogorov-Smirnov tests). TLR9^-/-^ mice were derived from 129/Ola x C57BL/6 F1 progeny [1] (http://www.myv.ne.jp/obs/index.files/tlr_eng.htm). b As for a using TLR9^M7Btlr^/Mmjax and 6J mice. TLR9^M7Btlr^/Mmjax mice have a TLR9 missense point mutation and do not respond to oligonucleotides containing CpG motifs (https://www.jax.org/strain/014534 ). c TLR9^-/-^ mice (like 6J) do not encode the full NNT gene. d Female 8-10 week old TLR9^-/-^ and 6J mice (n=6 mice and 12 feet per group) were infected with CHIKV and feet measured over time. Statistics by Kolmogorov-Smirnov tests. e Female C57BL/6J-Tlr9^M7Btlr^/Mmjax mice (n=6 mice and 12 feet per group) were infected as for d. f Viremia for the mice in d.
	After GzmA injection TLR9^-/-^ mice showed increased foot swelling (a), whereas C57BL/6J-Tlr9^M7Btlr^/Mmjax mice showed no significant difference (b). TLR9^-/-^ mice also have the NNT deletion (c); however, they are on a mixed 129/Ola and C57BL/6 background (Hemmi, H., et al., Nature, 2000. 408: 740-5), with 129/SvJ mice showing increased inflammatory infiltrates in certain settings (Hoover-Plow, J.L., et al., Inflamm Res, 2008. 57: 457-63). After CHIKV infection foot swelling was again increased in TLR9^-/-^ (d), but not TLR9^M7Btlr^/Mmjax mice (e). TLR9^-/-^ mice did not show an increased viremia (f). These data do not support a contention that TLR9 is required for GzmA’s bioactivity.
